# Novel Nuclear Roles for Testis-Specific ACTL7A and ACTL7B Supported by *In Vivo* Characterizations and AI Facilitated *In Silico* Mechanistic Modeling with Implications for Epigenetic Regulation in Spermiogenesis

**DOI:** 10.1101/2024.02.29.582797

**Authors:** Pierre Ferrer, Srijana Upadhyay, James J. Cai, Tracy M. Clement

**Affiliations:** Interdisciplinary Faculty of Toxicology Program, Texas A&M University, School of Veterinary Medicine and Biomedical Sciences, Texas A&M University, College Station, TX 77843; Department of Veterinary Physiology and Pharmacology, School of Veterinary Medicine and Biomedical Sciences, Texas A&M University, College Station, TX 77843; Department of Veterinary Integrative Biosciences, School of Veterinary Medicine and Biomedical Sciences, Texas A&M University, College Station, TX 77843

## Abstract

A mechanistic role for nuclear function of testis-specific actin related proteins (ARPs) is proposed here through contributions of ARP subunit swapping in canonical chromatin regulatory complexes. This is significant to our understanding of both mechanisms controlling regulation of spermiogenesis, and the expanding functional roles of the ARPs in cell biology. Among these roles, actins and ARPs are pivotal not only in cytoskeletal regulation, but also in intranuclear chromatin organization, influencing gene regulation and nucleosome remodeling. This study focuses on two testis-specific ARPs, ACTL7A and ACTL7B, exploring their intranuclear activities and broader implications utilizing combined *in vivo*, *in vitro*, and *in silico* approaches. ACTL7A and ACTL7B, previously associated with structural roles, are hypothesized here to serve in chromatin regulation during germline development. This study confirms the intranuclear presence of ACTL7B in spermatocytes and round spermatids, revealing a potential role in intranuclear processes, and identifies a putative nuclear localization sequence conserved across mammalian ACTL7B, indicating a potentially unique mode of nuclear transport which differs from conventional actin. Ablation of ACTL7B leads to varied transcriptional changes reported here. Additionally, in the absence of ACTL7A or ACTL7B there is a loss of intranuclear localization of HDAC1 and HDAC3, which are known regulators of epigenetic associated acetylation changes that in turn regulate gene expression. Thus, these HDACs are implicated as contributors to the aberrant gene expression observed in the KO mouse testis transcriptomic analysis. Furthermore, this study employed and confirmed the accuracy of *in silico* models to predict ARP interactions with Helicase-SANT-associated (HSA) domains, uncovering putative roles for testis-specific ARPs in nucleosome remodeling complexes. In these models, ACTL7A and ACTL7B were found capable of binding to INO80 and SWI/SNF nucleosome remodeler family members in a manner akin to nuclear actin and ACTL6A. These models thus implicate germline-specific ARP subunit swapping within chromatin regulatory complexes as a potential regulatory mechanism for chromatin and associated molecular machinery adaptations in nuclear reorganizations required during spermiogenesis. These results hold implications for male fertility and epigenetic programing in the male-germline that warrant significant future investigation. In summary, this study reveals that ACTL7A and ACTL7B play intranuclear gene regulation roles in male gametogenesis, adding to the multifaceted roles identified also spanning structural, acrosomal, and flagellar stability. ACTL7A and ACTL7B unique nuclear transport, impact on HDAC nuclear associations, impact on transcriptional processes, and proposed mechanism for involvement in nucleosome remodeling complexes supported by AI facilitated in silico modeling contribute to a more comprehensive understanding of the indispensable functions of ARPs broadly in cell biology, and specifically in male fertility.

## Introduction

Spermiogenesis represents one of the most drastic and complex cellular differentiation processes. The process includes significant restructuring of the nucleus impacting chromatin dynamics and transcriptional activity over the course of morphogenesis, with implications for epigenetic programing. The process of nuclear chromatin remodeling during spermiogenesis includes the replacement of the majority of canonical nucleosomes by protamines through a series of histone variant swapping and histone replacement steps. While these histone transitions have been well mapped over spermatogenic differentiation, the regulatory mechanisms along with potential implications for dynamic germ-line epigenetic programing remain to be fully understood. Given known roles of actin and actin related proteins in chromatin regulation of somatic cells, we here investigate the role of testis specific actin related proteins ACTL7A and ACTL7B as potential contributors to nuclear dynamics during spermiogenesis.

Actin is an omnipresent protein found in all known eukaryotes (Pollard, 2016). Over the billion years in which the actin protein family has been active, it comes as no surprise that it has diversified in function. Actins are known to be indispensable to life, and a crucial component in the orchestration of numerous and distinct cellular processes. The diverse functionality of actins is further compounded through their molecular divergence into further specialized Actin Related Proteins (ARPs). ARPs are also highly conserved proteins in eukaryotes, from yeast to mammals, sharing a common fold with actin. Some ARPs are known to contribute to filamentous polymers, such as ARP1 and ARP11 contributing to a filament in the dynein-dynactin complex, and ARP2 and ARP3 that contribute as branching and nucleating units in filamentous actin (Muller et al., 2005). However, ARPs have diverged from Actin in sequence and function, and have acquired even further specialized roles in distinct cellular contexts. ARPs often work collaboratively with actins in molecular processes including but not limited to signal transduction (Vasilev et al., 2021), programmed cell death (Ren et al., 2021), cellular locomotion (Verkhovsky et al., 1999), intracellular transport (Mori et al., 2011), and structural scaffolding (Vassilopoulos et al., 2019).

Amongst the most important and least understood roles of actins and ARPs is their involvement in intranuclear chromatin organization (Grummt, 2006; Baarlink et al., 2017; Xie et al., 2018). Like in the cytoplasm, actins exist in two forms in the nucleus: monomeric (G-actin) and filamentous (F-actin). Both the presence and polymerization state of nuclear actins and their concentrations are regulated by several factors, such as trans-nuclear trafficking, presence of actin nucleators, and actin sequestration by other Actin Binding Proteins (ABPs) (Kristó et al., 2016). From a macroscopic view of nuclear organization, actins, ARPs, and ABPs have been detected in several nuclear biomolecular aggregates including the nuclear dense lamina (Simon et al., 2010; Ranade et al., 2019), nucleoli (Katarína Kyselá et al., 2005; Castano et al., 2010), Cajal bodies (Skare et al., 2003; Gedge et al., 2005), nuclear speckles (Belin et al., 2013; Naum-Onganía et al., 2013), and transcription factories (Melnik et al., 2011). It is generally believed that in these nuclear suprastructures actin may provide structural support, scaffolding, or facilitate the movement or assembly of nuclear components. At large, actin has been demonstrated to be a crucial factor for gene translocation from the nuclear periphery to its association with nuclear speckles during gene induction (Khanna et al., 2014), and maintaining the 3D compartmental organization of the genome-wide architecture in mammalian cells (Mahmood et al., 2021).

In addition to the broader functionality of actins in nuclear organization, both actin and ARPs have crucial roles in more granular aspects of nuclear regulatory events pertaining to directed gene regulation and nucleosome remodeling. Actins are known to both interact with and affect the localization/recruitment of various nuclear proteins pertinent to gene expression and potential epigenetic encoding such as chromatin crosslinker HP1α (Toh et al., 2015), transcription factors PREP2 (Haller et al., 2004) and YY1 (Wu et al., 2007; Göös et al., 2022), transcription cofactors MRTFs (Gau & Partha Pratim Roy, 2018; Weston et al., 2012), RNA polymerases I-III (Philimonenko et al., 2004; Hofmann et al., 2004; Hu et al., 2004), Polycomb Repressive Complexes (PRCs) (Hauri et al., 2016), and nucleosome remodelers (Klages-Mundt et al., 2018). Of particular interest, ARPs are found to be stoichiometric components of several nucleosome remodeling complexes. These nucleosome remodeling complexes utilize ATP hydrolysis as an energetic cofactor to modify the structural composition or placement of nucleosomes through histone subunit exchange as well as sliding of the histone octamer across the DNA strand (Mazina & Vorobyeva, 2016). As a result, the activity of these remodeling complexes affects the accessibility of DNA to transcription factors and other regulatory proteins influencing gene expression. Two of the largest protein complex families responsible for altering histone octamer placement and composition in yeast and mammals alike are the INO80 and SWI/SNF families, respectively (Reyes et al., 2021). In addition to histone subunit swapping and nucleosome sliding, nucleosome remodeling complexes also include catalytic machinery required for the covalent addition or removal of Post Translational Modifications (PTMs) from the lysine-rich histone tails (Wang & Cole, 2020). The presence or absence of specific PTMs per histone subunit within a nucleosome often affects their structure but most importantly dictates their allowable interactions with other nuclear protein complexes, DNA, and/or higher-order chromatin structures. These modifications to the nucleosome are tightly regulated by enzymatic complexes containing specific members of discreet enzyme protein families such as histone acetyltransferases (HATs), histone deacetylases (HDACs), histone methyltransferases (HMTs), histone demethylases (HDMs), histone kinases, and histone phosphatases amongst many (Wang & Cole, 2020). In turn, the modification of histone subunits within a nucleosome lends itself as an “epigenetic code” to facilitate or discourage specific protein complexes from recognizing and thus interacting with the nucleosome (Waddington, 1942; Turner, 2007). The recognition of this epigenetic code is mediated through specific molecular domains contained within these protein complexes that in turn recognize specific PTMs such as histone acetylation by bromodomains; methylation by PHD Fingers, Tudor, and PWWP domains; and phosphorylation by BRCT, and BIR domains (Musselman et al., 2012). These histone PTMs frequently signal the recruitment of INO80 and SWI/SNF complexes. The opposite can also occur, where a preexisting nucleosome remodeler or the effects of one, such as the swapping of a histone subunit, in turn become the element which beckons PTM enzymatic machinery to the nucleosome (Wang & Cole, 2020; Demetriadou et al., 2020). Details regarding the intricate dance between different nucleosome remodelers and nucleosomes themselves are still poorly understood, but what has been observed as evident is the effect and presence of actins and ARPs on both sides of the equation.

One of the better understood mechanisms of how Actin and ARPs are recruited into and affect the function of nucleosome remodeling complexes is through their binding to a Helicase-SANT-associated (HSA) domain. As a long α-helical domain ∼75 amino acids long, the HSA domain was originally thought to be responsible for direct DNA binding but was later proven to be the primary binding site for ARPs and actin in both histone PTM enzymatic complexes and other nucleosome remodelers alike (Doerks et al., 2002; Horton et al., 2007; Knoll et al., 2018). Oftentimes the HSA domain within nucleosome remodeling complexes is found within the same subcomponent that houses the ATPase catalytic site. For example, mammalian SWI/SNF family members like neuronal BAF and PBAF complexes house their HSA domain in their catalytic subcomponents SMARCA2 and SMARCA4 both having their family’s conserved HSA amino acid sequence HQE(Y/F)LNSILQ (Eissenberg et al., 2005). Similarly, mammalian INO80 family members are also known to house their HSA domain in the same subcomponent as their ATPase catalytic site, with EP400 having both the ATPase catalytic site and HSA domain in the TIM60 (NuA4 mammalian equivalent) histone acetyltransferase complex (Fuchs et al., 2001; Elsesser et al., 2019) while SRCAP holds both ATPase and HSA domains in its complex (Eissenberg et al., 2005). Interestingly, but not surprisingly, INO80 family members have their own conserved HSA amino acid sequence HWDY(L/C)EEEM(Q/V) (Eissenberg et al., 2005) despite both INO80 and SWI/SNF HSA domains recruiting a remarkably similar subset of ARPs to an almost identical secondary structure within their respective complexes. The exact mechanism of how the binding of ARPs to HSA domains affects nucleosome remodeling function remains to be clarified. However, there is unambiguous evidence showing that the absence of specific HSA-binding ARPs in these complexes yields drastic changes to their overall functionality in terms of their ATPase activity (Szerlong et al., 2008), docking to the nucleosome (Yao et al., 2015; Ayala et al., 2018), and recruitment of additional cofactors like PTM enzymatic machinery and chaperone proteins (Percipalle, 2013).

The mechanistic directive of ARPs and nucleosome remodeling complexes have been better appreciated in unicellular organisms like yeast, with more recent progress in mammalian somatic cells. However, despite knowing that many of these molecular components are highly upregulated and present in germ-line meiotic and haploid cells, more is to be desired regarding knowledge of their general function in gametogenesis, the single most important cellular process regarding the genetic inheritance and propagation of all mammalian species.

Both ACTL7A and ACTL7B are testis-specific members of the larger ARP family and are required for male fertility. Our previous findings in knock out mouse models demonstrated that ACTL7A is required for subacrosomal F-actin formation and acrosome stability, and that ACTL7A is present in the intranuclear space of both spermatocytes and round spermatids (Ferrer et al., 2023). Furthermore, our characterization of ACTL7B as a key influencer of sperm formation and flagellar structural integrity included preliminary observations of nuclear localization (Clement et al., 2023; Ferrer et al, submitted). KO mice from both ACTL7A and ACTL7B also showed a drastic increase in DNA damage. A subjective coalition of these data showing DNA damage, compounding morphological defects, and intranuclear localization led us to speculate that these testis-specific ARPs may also serve functional intranuclear roles similar to nuclear somatic ARPs. This is of significant interest as testis-specific ARP contributions to nucleosome remodeling complexes could have implications for unique interactions or adaptive functions in germline development and epigenetics. Therefore, in this study we expand upon our previous findings by exploring regulation of ACTL7A and ACTL7B nuclear trafficking, the impact of ablation on gene regulation, histone and HDAC localizations during spermiogenesis, and further explore AI facilitated predictive models of ACTL7A and ACTL7B incorporation into chromatin regulatory complexes to propose detailed models for testis-specific ARP mechanisms in spermatid nuclear regulation.

## Results

### ACTL7B localizes to the intranuclear compartment of Pachytene spermatocytes and round spermatids

Our previous studies have demonstrated the intranuclear localization of ACTL7A in Spermatocytes and Spermatids (Ferrer et al., 2023) and speculated ACTL7B to have a potential intranuclear localization in spermatids (Clement et al., 2023; Ferrer et al. submitted). To further verify and characterize the intranuclear localization of ACTL7B, we now utilized a custom rabbit antibody specific to the n-terminus of ACTL7B (Clement et al., 2023) for refined assessment. Widefield fluorescent microscopy of ACTL7B immunofluorescence confirmed the previously characterized enriched acrosomal associated localization of ACTL7B (Clement et al., 2023(Fig. 1A). Interestingly, with the immunofluorescence staining protocol with this new antibody we also noted ACTL7B localization in spermatocytes, with enriched expression in a pattern consistent with synapsed chromosomal localization (Fig. 1B), as well as nuclear associated staining in round spermatids (Fig. 1A). To determine if this apparent nuclear staining observed with widefield microscopy was intranuclear or nuclear envelope-associated, we performed confocal microscopy line-scan analysis (Fig. 1C-E). We utilized the optical section with the largest nuclear diameter for each spermatid assessed to ensure line-scans through the intranuclear regions. We consistently observed intranuclear localization of ACTL7B in round spermatids through the initiation of elongation in step 8. Interestingly expression intensity in early round spermatid nuclei (Fig. 1C) was noticeably dimmer in intensity than those of late round spermatids (Fig. 1D-E), indicating increasing ACTL7B intranuclear localization as round spermatids progressively mature, consistent with a potential role in nuclear reorganization events during the acrosomal capping phase of spermiogenic progression.

**Figure 1:**
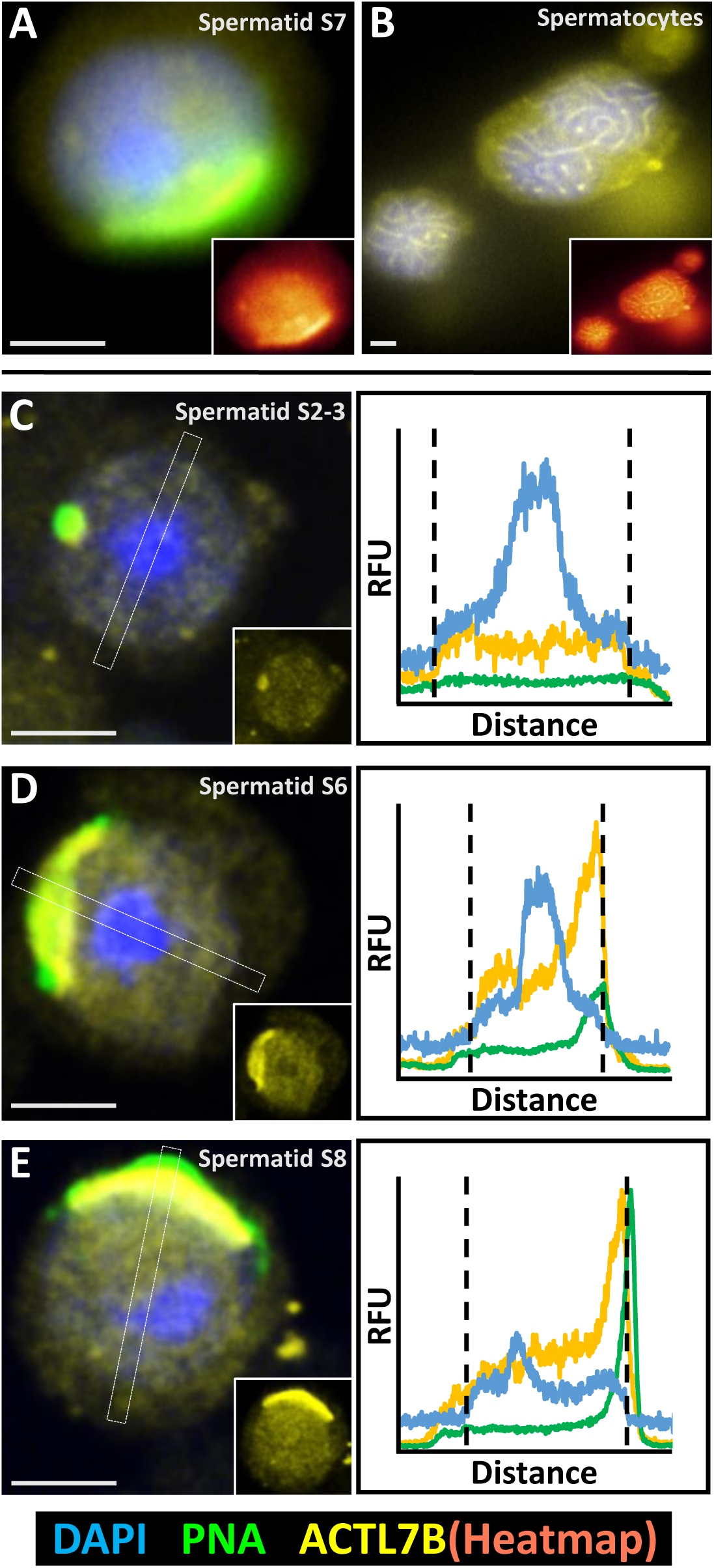
Native Intranuclear Localization of ACTL7B in Mouse Spermatocytes and Spermatids. (A-B) Widefield fluorescent microscopy showing intracellular localization of ACTL7B in spermatids (A) and spermatocytes (B) counterstained with DAPI and PNA for nuclear and acrosomal visualization. Subpanels show the fluorescent signal of ACTL7B as a heatmap gradient to better demarcate intracellular zones of ACTL7B enrichment. (C-E) Confocal microscopy images of round spermatids at different developmental stages, in the focal plane with the largest nuclear diameter selected for each cell to ensure intranuclear cross section. Localization of ACTL7B (yellow) is enriched in the nucleus (Blue=DAPI stained DNA) and subacrosomal space (Green= acrosomal PNA). Dashed rectangles in each image indicate the 25-pixel wide line-scan analysis area sampled in each cell to generate the signal plots to the right. Signal plots show the fluorescent intensity of each channel as Relative Fluorescent Units (RFU) across the distance (um) of each line scan. Vertical dashed lines in each signal plot demarcate the nuclear boundary within each cell. All white scalebars per image represent 2.5 um.

### ACTL7B has a functional NLS within its actin-like domain impacting nuclear affinity

Given the intranuclear localization of ACTL7B in the germline, we screened the sequence of ACTL7B across 42 mammalian species for conserved amino acid motifs characteristic of a Nuclear Localization Sequences (NLS). The criterion we utilized to identify a putative NLS included identifying highly conserved sequence segments which are Lysine/Arginine rich, have a basic isoelectric point, and are cytosolically accessible (i.e., being exposed at the protein’s surface) (Lu et al., 2021). Given the search parameters we were able to identify two potential NLSs within ACTL7B (Fig. 2), one being present near the end of the predicted N-terminal disordered domain between amino acids [K40-K50], and the other being a surface-facing α-helix within subdomain 4 of the actin-like body of ACTL7B between amino acids [H250-Y261].

**Figure 2:**
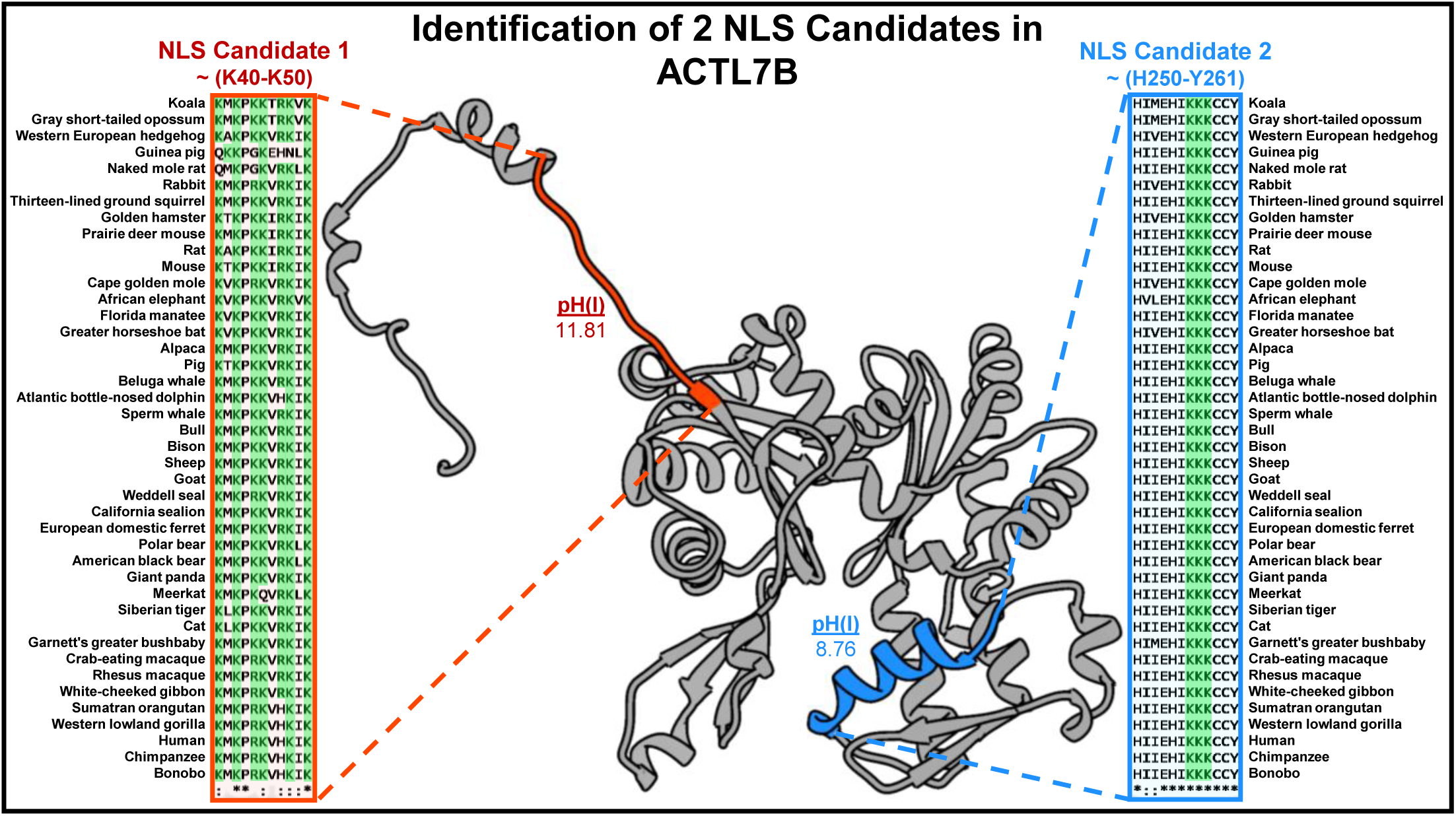
Highly Conserved Lys/Arg regions in ACTL7B Identified as Putative Nuclear Localization Signals (NLS). Two highly conserved Lys/Arg rich sequences were identified through multi-species sequence alignment of ACTL7B and hypothesized to be adequate nuclear localization signal (NLS) candidates. All lysine [K] and arginine [R] residues are highlighted in green for both alignment regions to illustrate sequence conservation and Lys/Arg enrichment regions. The regions of both NLS candidates are mapped onto the predicted structure of human ACTL7B with the color coordinated highlighting of residues showing NLS candidate 1 to be in the N-terminus domain (Red) of ACTL7B while NLS candidate 2 mapped to an external α-helix of subdomain 4 (Blue) of the actin body. Calculated isoelectric points, pH(I), are displaced for both sequences showing general basicity.

To evaluate the effectiveness of the two identified NLS candidates, we generated ACTL7B domain coding vectors along with positive and negative controls (Supplementary Fig. S1A-D) to assess the localization of protein products containing the putative NLS sequences in whole protein, protein subdomains, or independent NLS containing peptide sequences. Given that a candidate NLS was found in each of the two distinct and opposing domains within ACTL7B, we generated YFP-tagged truncated fusion protein vectors representing each of these two ACTL7B subdomains, which together represent the total protein sequence, and a whole ACTL7B-YFP fusion protein coding vector (YFP-7B). The truncated vectors coded for the actin-like domain [Leu56-Cys415] (YFP-7B(AD)), and the N-terminal domain of ACTL7B [Met1-Asp55] (YFP-7B(ND)) respectively. Additionally, we generated a YFP only coding vector as a negative control. Vectors were expressed in HEK293F (HEK) cells cultured in suspension. After determining the optimal post-transfection window to be 17-24 hours allowing for observation of localization patterns in cells balancing sufficient expression and maintaining higher viability (Supplementary Fig. 2A-B), we initially assessed localization of overexpressed full length YFP-7B. Assessment of YFP signal revealed full-length ACTL7B associates exclusively with the nucleus of the HEK cells (Fig. 3A), where this nuclear associated localization appeared mottled rather than equally diffuse. Of note, this mottled nuclear expression pattern for ACTL7B expressing HEK cells was consistently observed and did not exhibit the three specific differential nuclear associated localization patterns previously observed in ACTL7A transfected HEK cells (Ferrer et al., 2023). We next assessed localization of the actin and N-terminal sub-domains. To our surprise, the YFP-7B(AD) transfected HEK cells expressing the actin domain of ACTL7B phenocopied the nuclear affinity of full length ACTL7B (Fig. 3B) while YFP-7B(ND) expressing cells had the same cytosolic expression pattern as our YFP negative control (Fig. 3C-D). This result implicates the actin domain as responsible for the nuclear localization of ACTL7B, and not the disordered N-terminal domain. This finding was surprising since the N-terminus was considered to contain a better NLS candidate due to greater enrichment of Lys/Arg residues as well as the domain being substantially more flexible and cytosolically available given its disordered nature (Fig. 2). Furthermore, most somatic actins and ARPs are transported into the nucleus by dimerizing with Cofilin, and Cofilin’s NLS is then recognized by Importins and trafficked together with the associated actin to the nucleus (Kristó et al., 2016). Thus, it is entirely possible that the actin domain of ACTL7B is utilizing a similar transport mechanism for its nuclear localization rather than utilizing the ACTL7B encoded candidate NLS we identified. To more specifically evaluate the ability of the proposed NLS within the nuclear localizing ACTL7B actin domain to be recognized for nuclear transport, we generated two additional plasmids (Supplementary Fig. S1E-F). One plasmid expressed YFP conjugated to a small segment of murine ACTL7B [E242-A266] encompassing the second NLS candidate [EAGHKFSDDHLHIIEHIKKKCCYAA]. As a positive control, we used the same pcDNA6.2 backbone to express a well-known and characterized viral NLS [PKKKRKV] belonging to Simian Polyomavirus 40 (SV40) (Kalderon et al., 1984) also tagged with YFP for visualization. Transfection of our positive control SV40 NLS-YFP fusion protein product in HEK cells showed the expected YFP signal coming from the nucleus (Fig. 4A-B). In a similar fashion, the proposed ACTL7B NLS peptide sequence from the actin domain managed to transport the majority of the YFP fusion protein to the intranuclear region as seen by the enriched YFP fluorescent signal co-localizing with DAPI (Fig. 4C-D). The nuclear localization of the majority of the expressed YFP-7B NLS strongly suggests this 7B NLS sequence is sufficient for intranuclear transport and supports that a directly encoded NLS contributes to the observed intranuclear presence of ACTL7B in the germline. Interestingly, we do not observe the same subnuclear mottled patterning seen with whole ACTL7B or actin domain localization in HEK cells (Fig. 3A and B). The NLS peptide instead exhibits homogeneous distribution within the nucleus, indicating both a contribution in facilitating ACTL7B to the nuclear compartment, but also indicating that the peptide sequence alone is not sufficient for specific observed intranuclear associations once ACTL7B enters the nuclear compartment.

**Figure 3:**
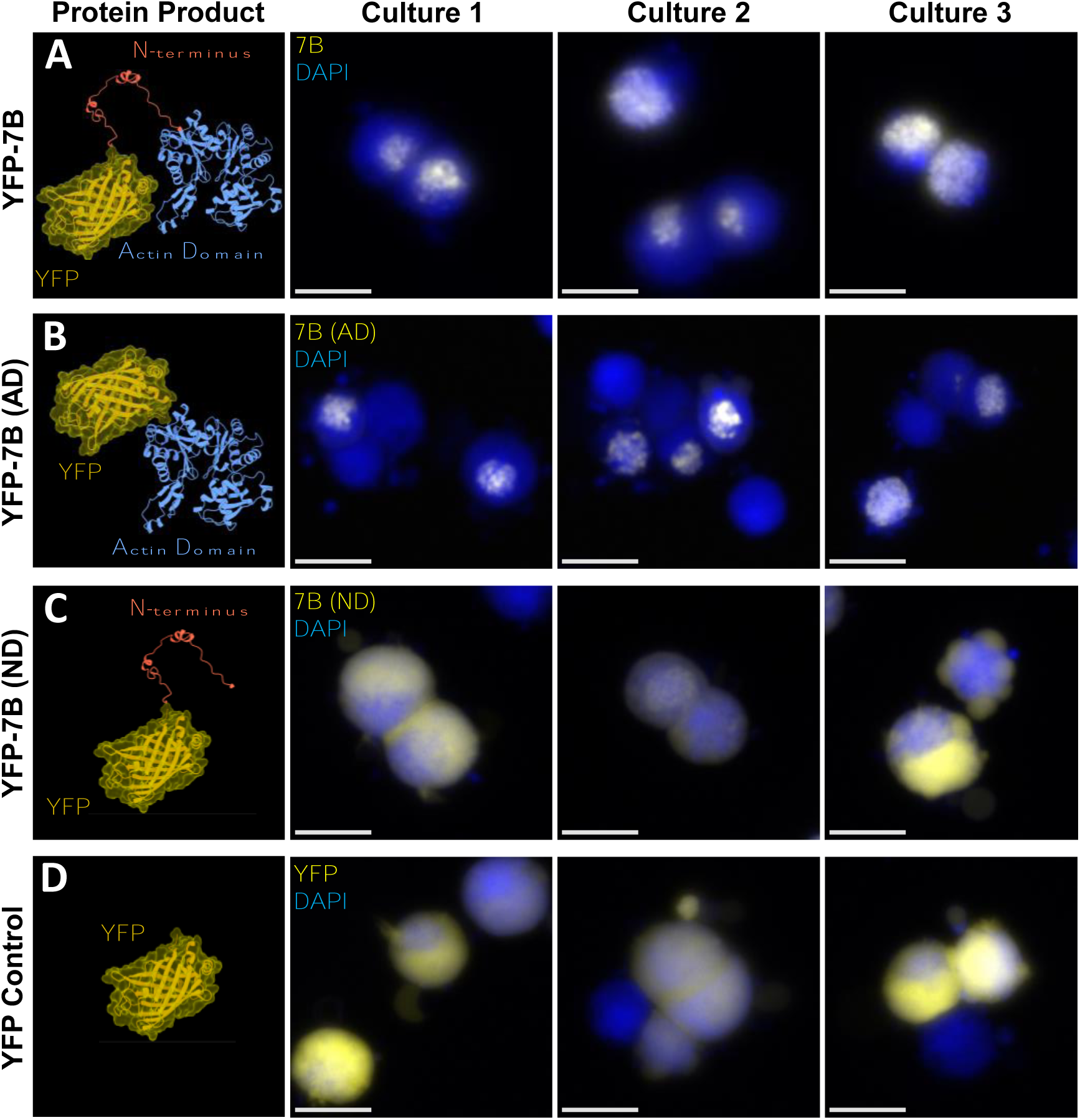
The Actin Body of ACTL7B is Responsible for its Nuclear Affinity. (A-B) Suspension culture HEK293F cells transfected to express YFP conjugated full length human ACTL7B (A), and YFP conjugated to the actin body only of human ACTL7B (B). Both full length and the actin domain of ACTL7B show a strong nuclear affinity as illustrated by the innate fluorescence co-localization of their conjugated YFP (yellow) with DAPI stained nuclei (blue). (C-D) Suspension culture HEK293F cells transfected to express YFP conjugated to the N-terminal domain only of human ACTL7B (C), or YFP alone as a negative control (D). The YFP control and YFP conjugated ACTL7B N-terminus expression patterns show similar cytoplasmic localization indicating no observable NLS activity from the over-expressed N-terminus of ACTL7B. Images are representative of triplicate cultures for all conditions. All white scalebars per image represent 10 ums.

**Figure 4:**
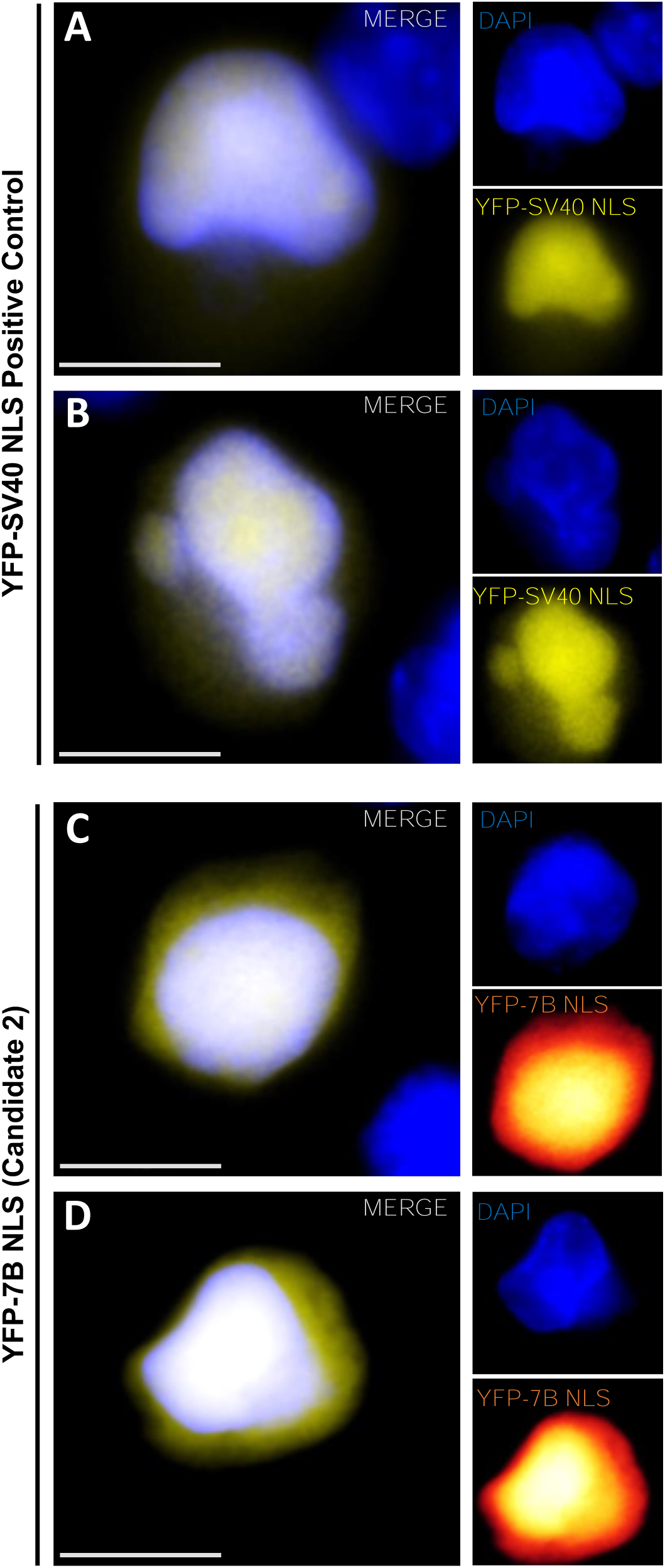
ACTL7B NLS Candidate 2 Induces Intranuclear Localization inHEK293F Cells. (A-B) Suspension culture HEK293F cells transfected to express YFP conjugated to the NLS of Simian Polyomavirus 40 antigen T [PKKKRKV] (SV40) as a positive control for intranuclear localization. Lateral subpanels deconstruct the merged image to its individual channels showing clear nuclear localization of the YFP fusion protein product. (C-D) Suspension culture HEK293F cells transfected to express YFP conjugated with a 25 residue peptide containing the second NLS candidate of ACTL7B [242-266]. The fluorescent signal of each channel is individually displayed as lateral panels with the YFP-7B NLS channel colored as a heatmap gradient showing both transient cytoplasmic expression and a highly enriched intranuclear signal.

### Loss of *Actl7b* leads to transcriptional upregulation of protease inhibition and immune/inflammation response related transcripts

Considering the known roles of other intranuclear ARPs in gene regulation, and the now clear presence of testis specific ACTL7B in the nucleoplasm, we postulated that loss of ACTL7B could impact gene expression within the testis. To investigate if there were any transcriptional consequences of ACTL7B ablation, we performed RNA-seq on whole testis samples isolated from 3 WT and 3 *Actl7b* -/- mice. Upon applying DESeq2 for data processing, we detected 385 transcripts exhibiting ≥ 2-fold upregulation in *Actl7b* -/- testes relative to wild-type tissue, whereas only 8 transcripts showed ≥ 2-fold downregulation (Fig. 5A). The predominantly upregulated transcripts observed, those with the largest fold change to p-value ratio, cluster into one or more of the following functional categories: 1) Genes expressing potent irreversible protease inhibitors like Serpina3n, Serping1, Serpina3g, and Serpina3i, as well as matrix-associated metalloproteinase inhibitor Timp1. 2) Genes driving cell death mechanisms like ferroptosis-associated genes Lcn2 and Tmem164, as well autophagy drivers like Hspb8 and complement pathway components C3 and C7. 3) Genes involved in immune and inflammation response including inflammatory caspase Casp4, innate immune initiator Masp1, extracellular apoptosis and inflammation nucleotide receptor P2ry2, interleukin receptors Il1r1 and Il17ra, and macrophage indicator Mpeg1. 4) Genes regulating cell adhesion and the extracellular matrix (ECM) such as cell-cell/matrix anchoring component Adam23, homophilic cell adhesion mediator Mpzl2, integrin-actin-tensin complex dissociation mediator Tns3, SLIT protein receptor Robo4, ECM cathepsin Cstb, and F-actin stress fiber disassembler Sema5a (Fig. 5A).

**Figure 5:**
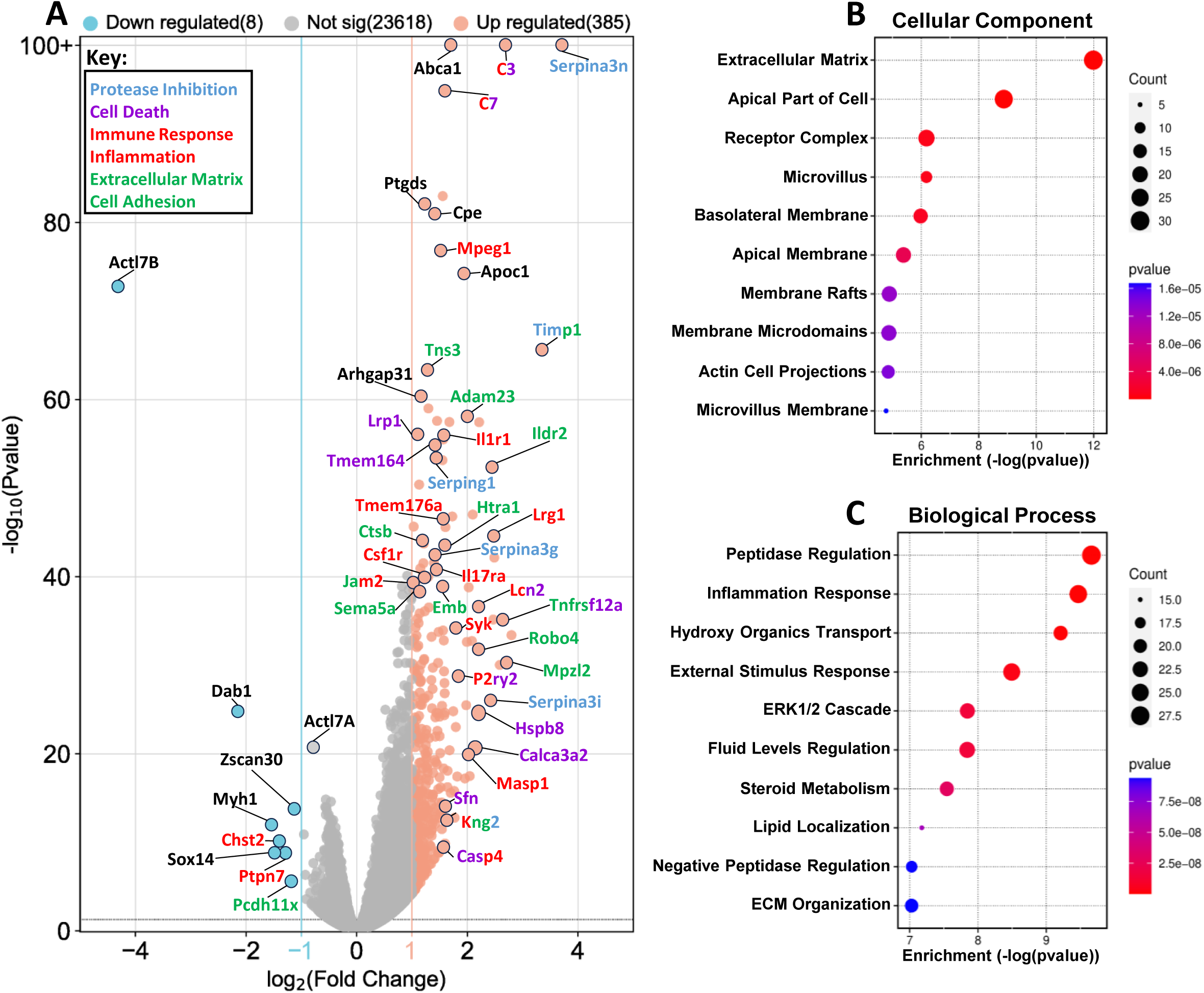
*Actl7B* Ablation Includes Gross Transcriptomic Consequences Including Altered Transcripts in Protease Inhibition, Inflammation, and Immune Response. **(A)** Volcano plot illustrating the transcriptomic differences between WT and *Actl7b -/-* murine testes. Fold change (FC) cut-offs are indicated as vertical lines at the -1 Log2(FC) position for downregulated genes (light blue) and the +1 Log2(FC) position for upregulated genes (salmon pink). Altered genes belonging to specific cellular mechanisms are color coordinated based on the provided key. (B-C) Gene ontology pathway analysis charts for cellular components (B) and biological processes (C) indicate unique categories for the altered genes between genotypes indicating gene count as circle size and significance of enrichment (p-value) as circle color ranging from a gradient scale of blue to red.

Gene Ontology assessment for cellular components and biological processes of the same data showed a similar trend, specifically indicating an increase in gene enrichment associated with the extracellular matrix, receptor and membrane complexes, peptidase regulation, inflammation response, and ERK1/2 pathways (Fig. 5B-C). The categorical congruity between these qualitative assessments leads us to conclude that loss of ACTL7B might affect not only the post-mitotic germline where it is expressed, but the testis as a whole through indirect mechanisms, thus implicating previously uncharacterized indirect contributions to the known infertility phenotype caused by ACTL7B ablation.

### Loss of *Actl7b* contributes to transcriptional changes of general and germline-specific transcription factors and regulators

Aside from the overt categorical transcriptional changes detected in the testis of *Actl7b -/-* mice, we observed additional significantly altered changes in transcription patterns between the WT and KO samples not recognized in the Gene Ontology assessments. Arrangement of the top 10% (2500) of transcripts with the highest absolute fold-change across biological replicates showed general transcriptional consistency between mice within the same genotype, illustrating a low level of inter-mouse variability within groups of biological replicates, thus improving confidence in the data and assessments (Fig. 6A). Processing this data through the TFCheckpoint 2.0 database (Chawla et al., 2013) we identified 47 known transcription factors (TFs) that were significantly altered past the ≥ 2-fold change threshold (45 upregulated and 2 downregulated) between WT and *Actl7b -/-* testis (Fig. 6B). Moreover, since our original query concerned the effects and roles of ACTL7B in gene regulation of the germline, we specifically looked at previously identified TFs uniquely expressed in different developmental stages of the male germline (Green et al., 2018) to determine if TF abundance might have been altered in the absence of ACTL7B. Through this approach, we detected that in *Actl7b -/-* testis, spermatogonia (SG) enriched TFs were mostly upregulated, preleptotene (PL) TFs were not appreciatively changed, spermatocyte (SC) and round spermatid (RS) TFs were moderately downregulated, and elongated spermatid (ES) TFs were more drastically downregulated (Fig. 6C). Overall, we observed an appreciable relationship between declining abundance of germline-specific transcription factors and spermatogenic progression in KO testis into phases where ACTL7B ablation would be expected to have higher impact, thus providing further support for the potential involvement of ACTL7B in germline gene regulatory events.

**Figure 6:**
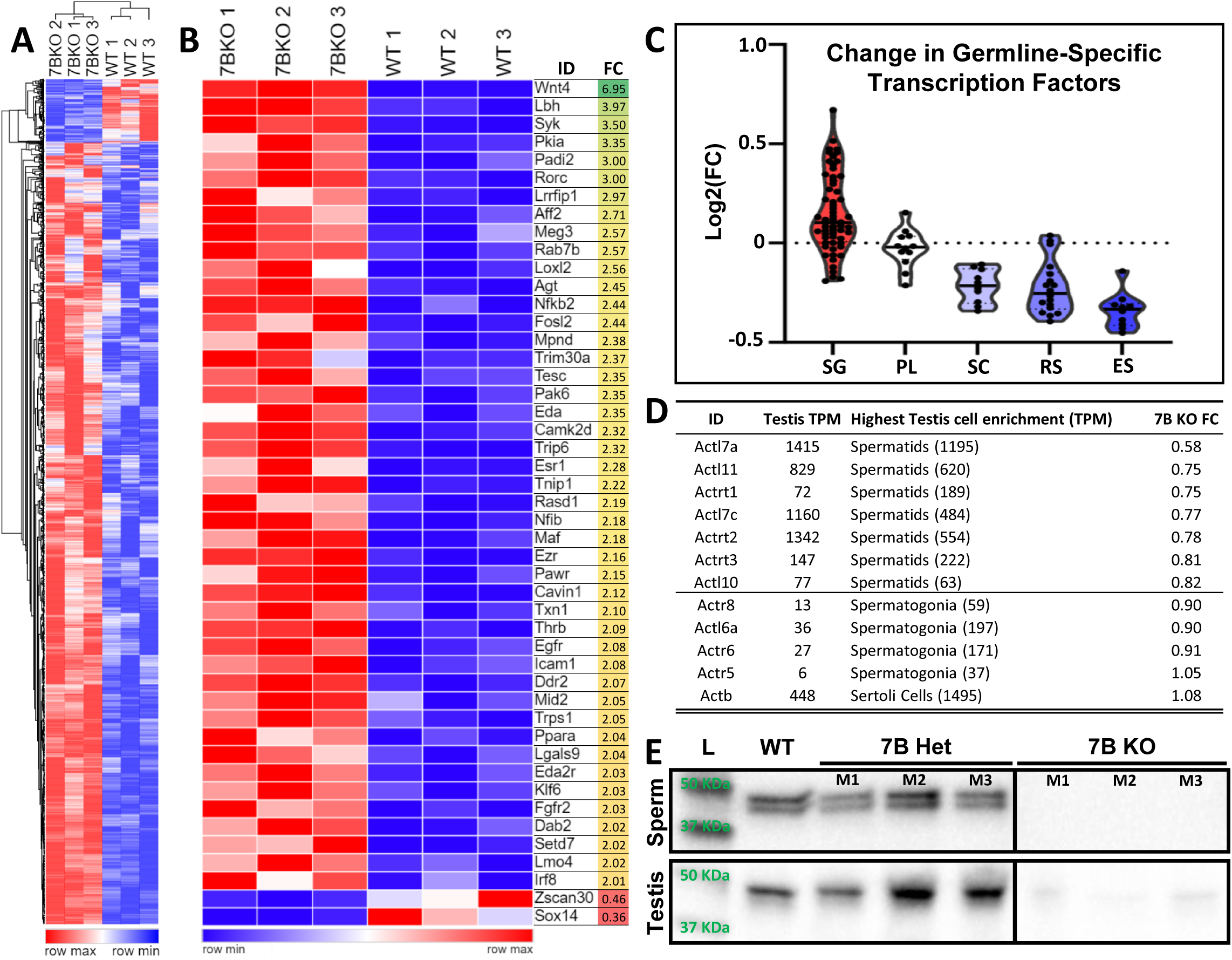
Loss of *Actl7B* Leads to Transcriptional Changes of General and Germline-Specific Transcription Factors and ARPs. (A) Heatmap of the top 2500 genes with the highest absolute fold change (FC) between WT and *Actl7b -/-* murine testes showing general transcriptional homogeneity between individuals of the same genotype. (B) Heatmap of all transcription factors with an absolute FC greater than 2 that were differentially expressed between genotypes. Heatmap is arranged in decreasing order of total FC per transcription factor. (C) Violin plots illustrating a trend in decreasing expression oftranscription factors previously identified as germ-line specific (Green et al., 2018) across developmentally progressive spermatogenic cell types caused by the absence of *Actl7b*; spermatogonia (SG); preleptotene (PL); spermatocyte (SC); round spermatids (RS); and elongating spermatids (ES). (D) List of testis-specific/enriched ARP genes (Top), and somatic nucleosome remodeling ARP genes (bottom) indicating their expected transcriptional prevalence in transcripts per million (TPM) in testes, and most enriched male germ cell (Robertson et al., 2020), as well as their overall FC between WT and *Actl7b -/- testes.* (E) Anti-ACTL7A IgG labeled western blot of testicular and sperm lysates from one WT, three *Actl7b +/-* (7B Het), and three *Actl7b -/-* (7B KO) mice indicating a severe reduction of ACTL7A expression in *Actl7b -/-* mice.

A conspicuous observation in the transcriptomic differences between *Actl7b -/-* and WT testis was the apparent downregulation of ACTL7A, reduced to 58% (FC 0.58) of its expected transcript prevalence compared to WT testis (Fig. 5A). Interestingly, in the absence of ACTL7B all other testis-specific ARPs had comparatively mild fold changes ranging from 0.82 to 0.75, while known somatic nucleosome remodeler-associated ARPs had little to no difference with fold changes between 1.08 and 0.9 in *Actl7b -/-* testis compared to WT (Fig. 6D). Given the observed transcriptomic changes in *Actl7a* -/- and our previous characterization of *Actl7a -/-* mice (Ferrer et al., 2023), we conducted western blot assessments on testes and sperm lysates from one WT, three *Actl7b -/+* (7B Het), and three *Actl7b -/-* (7B KO) mice to identify the effect of the aforementioned transcriptional downregulation of *Actl7a* on ACTL7A protein abundance (Fig. 6E). WT and 7B Het mice contained the expected band size with similar relative abundance for ACTL7A in both sperm and testis lysates. However, 7B KO mice contained only a faint band for ACTL7A in testis, and lacked an ACTL7A positive band in sperm (Fig. 6E). The decreased intensity in the ACTL7A signal in testis lysate from 7B KO mice is consistent with *Actl7a* transcriptomic reduction in the 7B KO testis. Furthermore, in sperm of the cauda epididymis we observe an apparent loss of ACTL7A retention in sperm from 7B KO mice. It is possible that ACTL7A is still present in the sperm of 7B KO sperm at low levels insufficient to be detected in standard western blotting practices. Alternatively, it may be that in the absence of ACTL7B the ACTL7A remaining in 7B KO spermatids is not properly incorporated or stabilized into the postacrosomal sheath where ACTL7A has been shown to reside in sperm (Ferrer et al., 2023). Additionally, any ACTL7A still present in 7B KO sperm could be lost during epididymal transit or during cell isolation procedures given the extreme morphological deformities and general fragility that 7B KO sperm portray (Clement et al., 2023). Overall, the specific changes in general and germline-specific transcription factors, as well as reduction in *Actl7a* transcription and translation in 7B KO testis is indicative that *Actl7b* ablation can have resounding gene regulatory consequences supportive of potential involvement in such mechanisms as is known for somatic expressed nuclear ARPs.

### *In silico* modeling of ACTL7A and ACTL7B interactions supports the capacity to bind the HSA region of nucleosome remodelers

Given the evidence presented here for ACTL7A and ACTL7B nuclear localization and impact on gene regulation, we sought to test if these testis-specific ARPs could participate in conjunction with, or in place of, known intranuclear chromatin remodeler associating ARPs such as ACTL6A, ACTL6B, ACTB or ARP6 using an *in silico* modeling approach. ARPs are known to directly contribute to the function of nucleosome remodelers conserved from yeast to mammalian somatic cells through two salient yet distinct mechanisms. In one chromatin remodeling complex interaction, ARPs serve as a stabilizing buttress between the rest of the chromatin remodeling protein complex and associated nucleosome, as is the case with ARP6 and the human SRCAP complex (Feng et al., 2018), as well as ARP5 with the INO80 complex in yeast (Zhang et al., 2023). The second conserved interaction in which ARPs associate in chromatin remodeling complexes is more amorphous, where one or more consecutive ARPs or Actin monomers bind to the HSA domain within the nucleosome remodeling complexes. A large multi-protein component of one such nucleosome remodeling complex, the SMARCA4 PBAF complex, has recently been structurally elucidated via Cryo EM for humans (Yuan et al., 2022). Through structural and interface analysis of the published structure via ChimeraX (PDB: 7VDV), we have illustrated the spatial relationship between the different molecular components of this partial complex (Fig. 7A-C). Here, one can appreciate the intricacies of the nucleosome-bound PBAF complex, such as how it is evident that SMARCA4 (represented in light and dark orange, Fig. 7A) is the core scaffolding protein of this complex as it stabilizes and interacts with almost every other component of the PBAF complex including the ARP subunits, DNA strand, and histone octamer (Fig. 7A-B). However, it is important to note two specific details regarding ACTB and ACTL6A, the ARP and Actin monomers present in this complex. 1) These actins are in very close spatial proximity to the nucleosome (Fig. 7A) but surface interaction and interface mapping analyses from this cryo EM elucidated partial complex are not suggestive of direct interactions between ACTL6A or ACTB with either the DNA strand nor the histone octamer (Fig. 7B). 2) Despite the robust interactions between many of the protein components within the PBAF complex, ACTB and ACTL6A not only exclusively bind to the HSA domain of SMARCA4, but they do so in an adjunctive manner in which both actin monomers interact with one another (Fig. 7A-B). Binding of ARPs to the HSA domain of many nucleosome remodelers, as well as the adjunctive fashion in which such ARP and actin monomers interact with each other in the complexes, has been readily documented in yeast and mammals alike (Reyes et al., 2021; Bieluszewski et al., 2023). Additionally, many of these ARPs such as ACTL6A/B (Karlsson et al., 2021), as well as nucleosome remodelers such as SRCAP complex members (Karlsson et al., 2021; Sun et al., 2022), SMARCA family members (Chong et al., 2007; Farshad Niri et al. 2021), INO80 (Serber et al., 2016), and EP400 (Karlsson et al., 2021), have been documented to be present across many developmental stages of the male germline. With the provided evidence at hand holding ACTL7A and ACTL7B as candidates for engaging with germline expressed conserved nucleosome remodelers in a similar fashion as their not-so-distant protein relatives, we sought to test the feasibility of such molecular interactions beginning here with an *in silico* modeling approach

**Figure 7:**
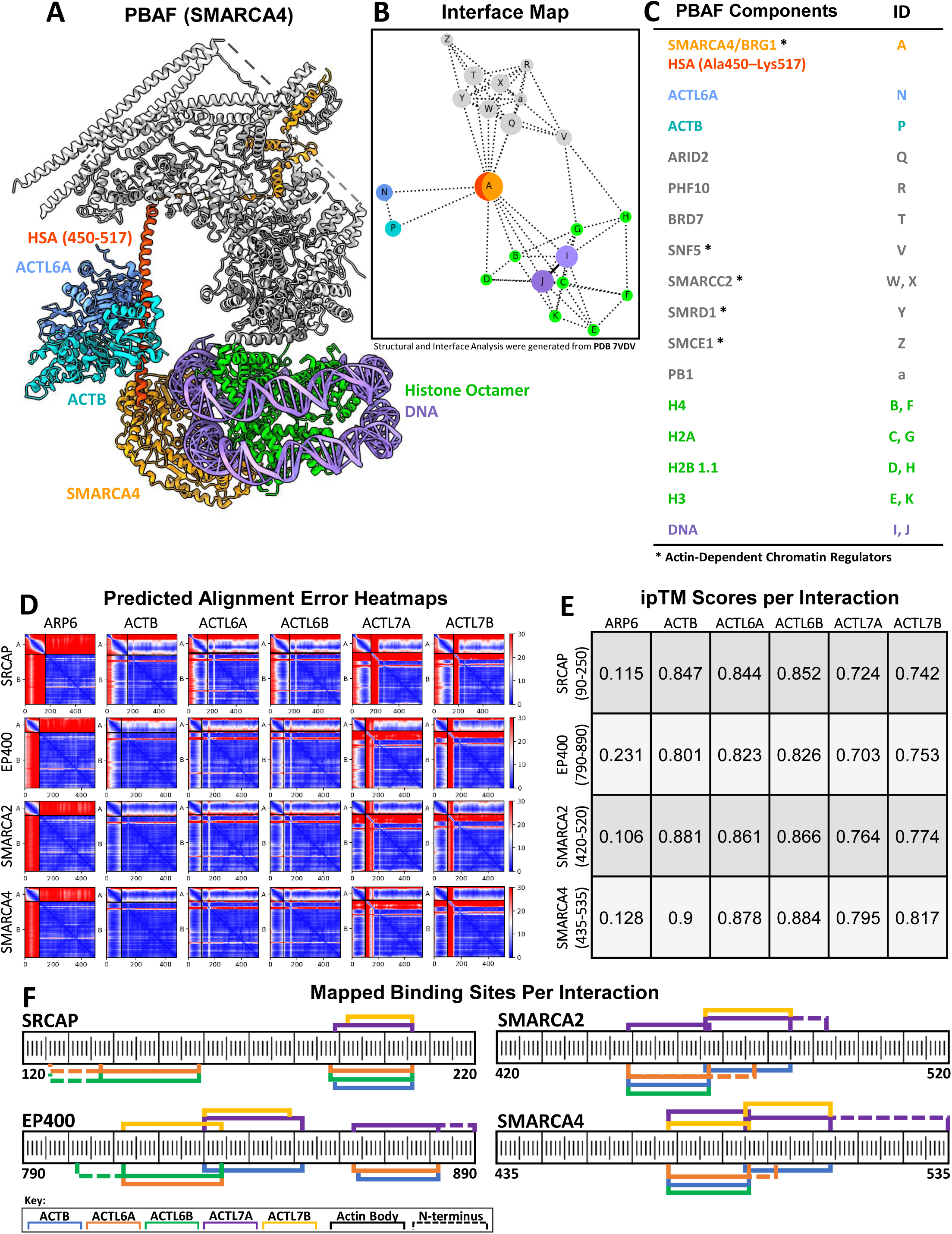
*In Silico* Modeling Indicates Positive Predictive Value for the Binding Capacity of Testis Specific ACTL7A and ACTL7B to HSA Domains of Classical Nucleosome Remodelers. (A-C) Generated structural model of human PBAF nucleosome remodeling complex (PDB: 7VDV) bound to ACTL6A and ACTB (A), with interface map analysis of protein interactions through ChimeraX (B), and list of protein components (C). (D) Predicted alignment error heatmaps of the top dimeric interactions between ARP6, ACTB, ACTL6A, ACTL6B, ACTL7A, and ACTL7B with the isolated HSA domain of SRCAP, EP400, SMARCA2, and SMARCA4. On each alignment error heatmap red indicates poor to no predicted binding, while blue represents predictive values indicating excellent binding potential between protein residues. The presence of a clear blue vertical strip at the interaction quadrant of each HSA domain with each ARP (bottom left quadrant of each heatmap) indicates strong binding affinity between the two proteins at those residues, absence of a blue line corresponds to poor to no binding affinity. (E) Tabulated results of the interface predicted template modeling (ipTM) score of each ARP’s binding stability with each HSA domain from a scale of 0 to 1.0. An ipTM score of 0.7 or higher for a dimeric prediction is seen as very favorable and biologically relevant. (F) Linear HSA domain primary protein sequence maps with colored brackets indicating all stable predicted ARP interactions (those with an ipTM score higher than 0.7) between each modeled HSA domain and the select ARPs. Amino acid sequence residue numbers corresponding to the parent chromatin remodeling protein are indicated below each represented HSA domain.

Here we sought to capitalize on the recent advancements in artificial intelligence driven structural predictive models, such as AlphaFold2 Multimer (Jumper et al., 2021; Evans et al., 2022), to test the feasibility of ACTL7A and ACTL7B binding to the HSA domains of select nucleosome remodelers in the germline, while simultaneously assessing the efficacy of these novel high-powered AI-driven tools to better contextualize predictive values for further research. Due to the novelty of the AlphaFold2 Multimer model, not a lot of evidence exists proving the efficacy and accuracy of its predictions regarding nucleosome remodeling complexes and their associated ARPs. To test the conformity of AlphaFold2 predictions to known biological structures and interactions of nucleosome remodelers we first set out to test if we could identify the HSA region of a nucleosome remodeler solely through its predicted interaction with one of its known binding partners to serve as a positive control for predicted interactions. Additionally, we wanted to make sure that the predictive model wouldn’t erroneously endorse a fictitious interaction between an HSA domain and an ARP that is known in previously published literature to interact with nucleosome remodelers but not via its HSA domain, thus serving as a negative control for predicted interactions in the HSA domain. To evaluate these control parameters, we chose the SRCAP chromatin remodeling protein complex. Given that SRCAP meets the criteria of having both an HSA binding ARP (ACTL6A) and a non-HSA binding ARP (ARP6) within its multimeric complex (Feng et al., 2018), as well as having a sufficiently large body of published investigative literature, we chose it as an appropriate target to test the AlphaFold2 *in silico* predictive model and validate predictive values relevant for alternative ARP subunit binding.

Initial modeling of the well-established interaction between the SRCAP HSA domain [M1-K706] and ACTL6A resulted in an extremely high-fidelity predicted interaction with an ipTM score of 0.853 out of 1.0. This allowed confident identification of ACTL6A binding to the specific HSA region of SRCAP between amino acids (AA) 90 and 250 (Supplementary Fig. S3A-B), thus matching known biologically relevant associations (Yin et al., 2022; Evans et al., 2022). Additionally, the *in silico* model did not predict any biologically aberrant interactions between non-relevant SRCAP residues outside the HSA domain and ACTL6A. Lastly, models predicting the interactions between ARP6 and SRCAP [M1-K706] showed minimal stability between the two proteins, having an ipTM score of 0.237, appropriately indicating low potential for biological relevance of the predicated interaction of the ARP6 in the ARP binding region of the SRCAP HSA domain (Supplementary Fig. S3C-D). The results of these structurally evidenced benchmarked controls affirmed confidence in the predictive capabilities of AlphaFold2 for biological relevance of ARP-HSA interactions. We proceed with using the predictive model to test the interactions of ACTL7A and ACTL7B with isolated HSA domain containing fragments of two INO80 family members, SRCAP [E90-S250] and EP400 [K790-I890], as well as two SWI/SNF family members, SMARCA2 [R420-Q520] and SMARCA4 [N435-E535] (Fig. 7D). Aside from the predictive interaction modeling between ACTL7A and ACTL7B and the select HSA domains, we also ran predictive interaction modeling of those same HSA domains with three known positive controls from the literature, ACTL6A, ACTL6B, and ACTB (Knoll et al., 2018; Klages-Mundt et al., 2018) and the previously mentioned negative control ARP6 (Fig. 7D).

Tabulating the ipTM results of all assessed interactions illustrated a very clear pattern in the predicted strength of each interaction between the select ARPs and HSA domains. ARP6 proved to be a faithful negative control; despite being a close family member to the other ARPs, it had a low ipTM score range between 0.106 and 0.231 across HSA interactions (Fig. 7E). Similarly, ACTB, ACTL6A, and ACTL6B were reliably consistent as positive controls in providing interactions with ipTM scores exceeding that required to indicate biologically relevant association with a range between 0.801 and 0.9 (Fig. 7E). The experimental predictions of interest between ACTL7A or ACTL7B with each of the four chosen HSA domains did not reach an average level of interaction strength as high as the known somatic cell binding positive control ARPs. However, their ipTM scores did surpass thresholds indictive of biological relevance with a score range between 0.703 and 0.817 (Fig. 7E), resulting in the interpretation that ACTL7A and ACTL7B could serve as testis specific ARP components within nucleosome remodeling complexes in the germline and thus have the potential to impact chromatin and gene regulation in the male germline through mechanisms conserved across species for somatic nuclear ARP paralogs.

In addition to quantifying the strength of each paired interaction to represent potential for biologically relevant interactions, we also analyzed the generated PDB files of each predicted interaction using ChimeraX to map and better understand the positions each ARP could linearly occupy within each HSA domain (Fig. 7F). Visualizing the mapped binding sites of each ARP across HSA domains revealed some interesting patterns in the way the ARPs behave between the two INO80 and SWI/SNF nucleosome remodeler families. Initially, it became apparent that SMARCA2 and SMARCA4 had a very similar ARP specific binding localization patterns. Likewise, SRCAP and EP400 more closely resembled each other in their ARP specific binding patterns, but not as similarly as the comparison between SMARCA4 and SMARCA2 (Fig. 7F). This result was somewhat expected given that the shared similarity in ARP binding patterns between the HSA domains correlates with nucleosome families, given that SMARCA2 and SMARCA4 are both SWI/SNF nucleosome remodelers while SRCAP and EP400 both belong to the INO80 family. Remarkably, the spatial pattern in ARP binding sites of both SRCAP and EP400 is highly suggestive that three or even four ARP and/or actin monomers could bind sequentially to those respective HSA domains (Fig. 7F). This observation has been structurally confirmed in the yeast INO80 complex, with the adjunct monomers of Arp4, Actin, and ARP8 all binding in tandem to the INO80 HSA domain (Knoll et al., 2018; Zhang et al., 2023), but similar tandem multi-ARP associations have yet to be structurally confirmed in humans. Our predictive models for ARP binding of SMARCA2 and SMARCA4 HSA domains suggest a maximum of two ARP and/or actin subunits would be likely to bind these SWI/SNF remodeler HSA domains within the two adjacent binding sites predicted between all modeled associations (Fig. 7F). An interesting observation that can be made with the ARP binding patterns across SMARCA2 and SMARCA4 is, that unlike ACTL6A or ACTL6B, both ACTL7A and ACTL7B are predicted to be able to replace ACTB in its known binding regions of each HSA respectively (Fig. 7F). For example, this would indicate that a PBAF complex found in the germline could have the structurally established ACTL6A-ACTB arrangement docking to its HSA domain, or a now theoretical ACTL6A-(ACTL7A/ACTL7B) conformation which could alter the specificity and function of the PBAF complex in a manner unique to the germline.

### *In silico* models illustrate that ACTL7A and ACTL7B can adopt varying binding conformations relative to ACTL6A when cooperatively binding the HSA domain of SRCAP

We next used these predictive tools to investigate how multiple ARP monomers could disrupt or cooperatively bind to a single HSA domain. This was an aspect we considered to be more pertinent to the actual function and relevance of these complexes in a biological system. Owing to the observed reliability of SRCAP as a model protein with our previous control testing (Supplementary Fig. S3) and its expected allowance of multiple ARPs to cooperatively bind its HSA domain (Fig. 7F), we tested if we could predict any adjunctive behavior between the ARPs when binding an HSA using this model. Predicted interactions between ACTL6A and the actin bodies of ACTL7A [A71-F435] and ACTL7B [A51-C415] on the HSA domain of SRCAP reaffirmed our previous speculation of SRCAP binding at least these three ARPs with strong predictive values for biological relevance (Fig. 8A-B). More interestingly, the predicted arrangements between ACTL6A, ACTL7A, and ACTL7B with the SRCAP HSA had some unexpected results. For example, both ACTL7A and ACTL7B were predicted to bind a novel region of the SRCAP HSA domain between [D150-H173] for ACTL7A (Fig. 8A) and [E154-H173] for ACTL7B (Fig. 8B). These novel conformations for ACTL7A and ACTL7B are both adjacent to and interact with ACTL6A as it sits on its previously predicted binding site with SRCAP [H136-R159] (Fig. 7F), (Fig. 8A-B). This result would indicate that both ACTL7A and ACTL7B require ACTL6A to be present to stabilize the interaction at the novel binding site within the HSA domain of SRCAP. Moreover, this result also opens to interpretation the idea that even with the same stochiometric ratios of different ARPs, their sequential binding locations across an HSA domain is not entirely deterministic as we have observed here with the predictive interactions between ACTL7A, ACTL7B, and ACTL6A with SRCAP (Fig. 8 A-B). Next, given the 14-18 amino acid gap in the HSA domain of SRCAP between ACTL7A and ACTL7B (Fig. 8A-B) we proceeded with testing if an additional ARP could bind this vacancy in a manner that further stabilized ACTL7A and ACTL7B. For this additional experiment we incorporated either ACTB or ACTL6B into the predictive space with ACTL6A, ACTL7A, ACTL7B, and the HSA domain of SRCAP. When adding ACTB into the fold, we consistently observed ACTB to supplant both ACTL7A and ACTL7B as the primary interactor of ACTL6A when bound to the HSA domain (Fig. 8C). Moreover, when ACTL6A and ACTB dimerized and bound to the HSA domain, ACTL7A was able to outcompete ACTL7B for their previously observed binding site at SRCAP [L191-W206] (Fig. 8C). In contrast, when ACTL6B was introduced into the system, we observed ACTL6A to exclusively bind with ACTL7B and stabilize its interaction with the HSA domain. In this model, ACTL6B and ACTL7A seemed to interact in a bipartite system in which ACTL6B bound ACTL7A but did not directly interact with the HSA, instead preferring a bent conformation of the HSA domain in the predicted association. Alternatively, in the presence of monomeric ACTL6B, ACTL7A failed to bind SRCAP (Fig. 8D). Since these two interactive states appear to potentially be endpoints of a systematic interaction, it is difficult to interpret weather ACTL6B may be recruiting ACTL7A to the HSA domain or if it was displacing an already HSA-bound ACTL7A to dissociate it from the complex. The latter interaction, in our opinion, seems more likely considering that ACTL7A readily bound the HSA domain of SRCAP without ACTL6B present (Fig. 8B) and ACTL6B never dimerized with ACTL7A in isolation in these models (i.e., when ACTL7A wasn’t already interacting with an HSA domain) (Fig. 8D).

**Figure 8:**
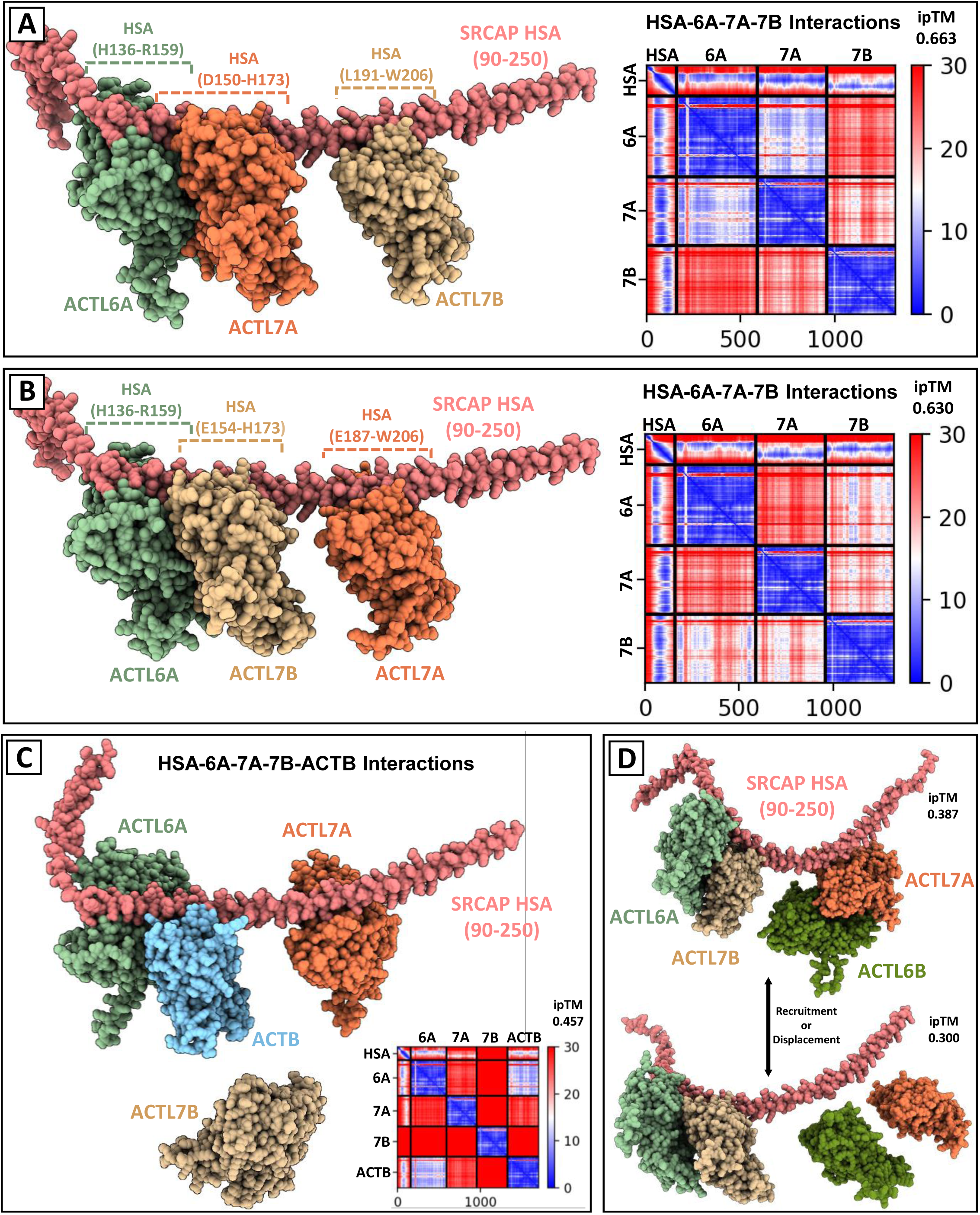
ACTL7A and ACTL7B Actin Domains Predicted to Interact with the HSA Domain of SRCAP Cooperatively with Classical Somatic ACTL6A. (A-B) cooperative binding model of ACTL7A and ACTL7B with ACTL6A and the HSA domain of SRCAP showing an ability of SRCAP to supportively bind at least three ARPs in tandem and in more than one conformation. Accompanying predictive alignment error heatmaps show the direct binding capabilities between these protein components. (C) Multi-ARP competitive binding model between ACTL6A, ACTL7A, ACTL7B, and ACTB with the HSA domain of SRCAP showing the predicted preferential occupancy of ACTB to the HSA binding site adjacent ACTL6A, thus supplanting ACTL7B in this conformation. (D) Additional competitive binding model with ACTL6B instead of ACTB indicating predicted potential to aid in the recruitment or displacement of ACTL7A from its binding site to the HSA domain of SRCAP .

When revisiting the predicted models between the HSA domain of SRCAP with ACTL6A, ACTL7A, and ACTL7B for further examination, we noticed two distinct structural features that could be pertinent regarding their function as ARPs in a nucleosome remodeling complex. Much like the ARP module of yeast INO80 (Knoll et al., 2018; Zhang et al., 2023), the HSA domain of SRCAP is unilaterally enriched with lysine/arginine residues on the face of the α-helix that would contact the DNA strand of an associated nucleosome (Fig. 9A, C). This by itself is not strange since lysine and arginine are notoriously well documented to strongly bind the poly-negative backbone of DNA and participate in nucleotide readout (Martin et al., 2022; Kazi Amirul Hossain et al., 2023). Additionally, many DNA binding proteins such as TATA-Box Binding Proteins (Pardo et al., 2000) and the HSA domain of yeast INO80 (Knoll et al., 2018) are known to interact with DNA strands through these amino acids. What makes this observation intriguing is that the base of the disordered N-terminal domains of both ACTL7A and ACTL7B are also enriched with lysine/arginine residues and are unequivocally positioned in the direction that would face a DNA strand that would interact with the lysine/arginine rich side of the HSA domain (Fig. 9A, C). This could potentially explain one of the possible functions of the identified lysine/arginine rich region in the N-terminus of ACTL7B which was previously thought to be an NLS candidate (Fig. 2). Additionally, we observed that the manner in which ACTL6A, ACTL7A, and ACTL7B bind to the HSA domain of SRCAP ends up orienting the pointed ends (subdomains 4 and 2) of each ARP away from the HSA domain. Such adopted conformation also results in the exposure of the DNase-binding loop (D-loop) of each bound ARP (Fig. 9B-C). The D-loop is a structural motif conserved within the second subdomain of many actins and ARPs (Muller et al., 2005; Pollard, 2016). It is mostly comprised of a continuous disordered loop terminated by a small antiparallel β-sheet formed by the start and end of this structure (Muller et al., 2005; Pollard, 2016). Although the structure of the D-loop between ARPs is similar, both their specific purpose and sequence identity can vary, resulting in ARP-specific convoluted functions. Known functions of the D-loop include the recruitment of DNase I (Kabsch et al., 1990), altering the catalytic activity of ATP hydrolase sites (Durer et al., 2012), varying the thermal stability of actins (Levitsky et al., 2008), facilitating F-actin branching (Chou et al., 2022), binding motor proteins (Kubota et al., 2009), and possibly recognizing specific histone variants (Saravanan et al., 2012). The illustrated structural models do not infer the specific functions of the D-loop of ACTL7A, ACTL7B, nor ACTL6A. However, these results indicating coordinated directional orientation are suggestive that they might play a spatial organizational role in additional subunit recruitment to a nucleosome remodeling complex given the varied interacting partners reported for these D-loop domains (Fig. 9B-C).

**Figure 9:**
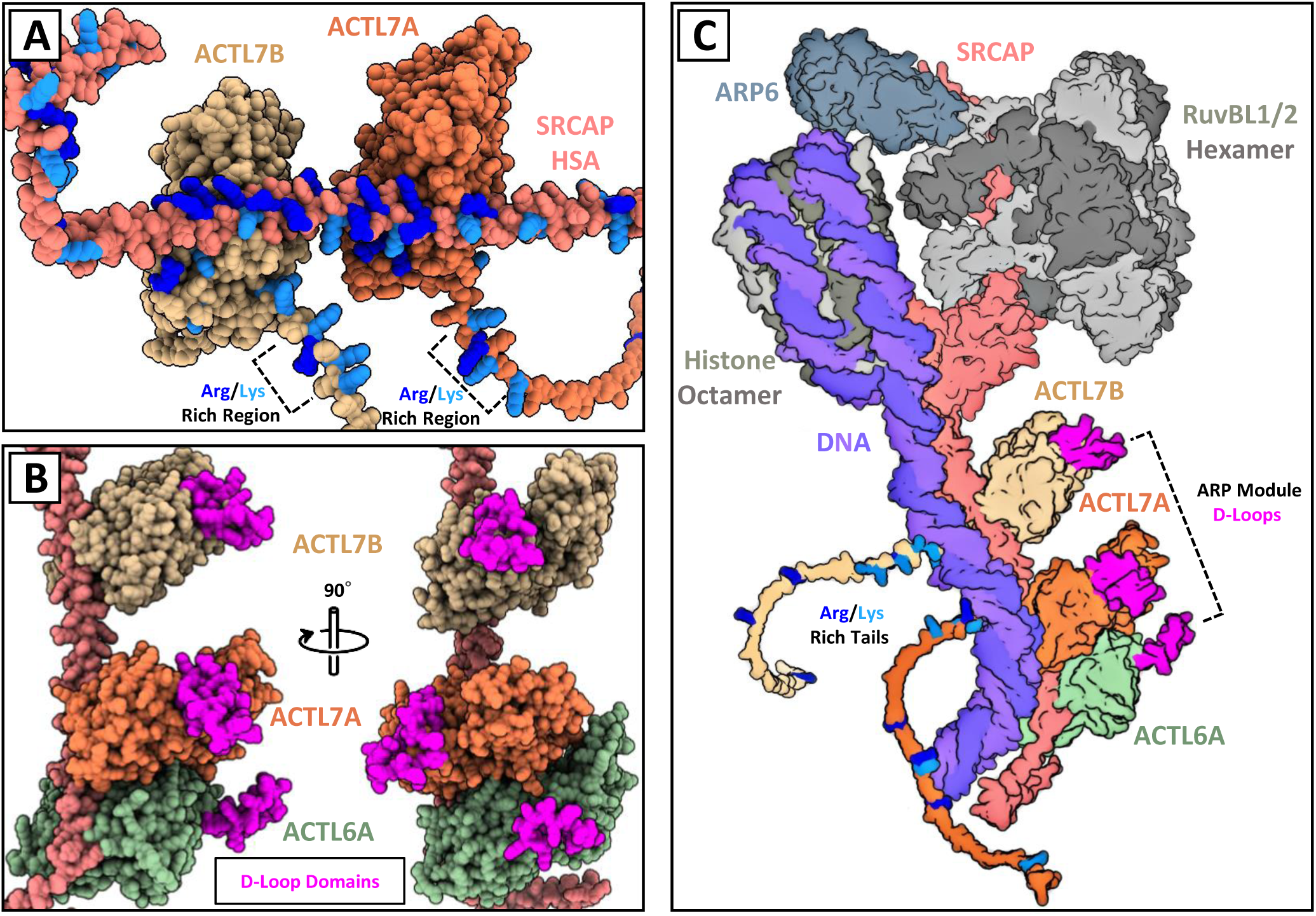
Full length ACTL7A and ACTL7B Orientation in a SRCAP Complex is Predicted to Poise N-terminal Disordered Domains for Possible DNA Interactions, and D-loop Domains for Accessory Protein Interactions. (A) Predictive interaction between ACTL7A, and ACTL7B with the HSA domain of SRCAP indicating a unilateral enrichment of Lys/Arg residues contained in the HSA domain and in the N-terminal domain of both ARPs indicating a potential special orientation between these components to bind DNA directly. (B) Predictive model of the binding between ACTL7A and ACTL7B with ACTL6A and the HSA domain of SRCAP showing a unilateral arrangement of the D-loop domains of all three ARPs involved. (C) Artistic schematic of the complete chromatin associated human SRCAP nucleosome remodeling complex to illustrate the proposed interaction model contextualizing how testis specific ACTL7A, ACTL7B could structurally integrate with classical somatic ACTL6A and SRCAPto form a putative testis specific regulatory ARP module. This schematic was generated by substituting the recently elucidated SRCAP partial complex (PDB: 6IGM) where the known yeast INO80-RuvB hexamer complex would be relative to its ARP8-Actin-ARP4-HSA-DNA Arp module, and bound nucleosome, as proposed by (Zhang et al. 2022). We then substituted our predictive ACTL7A-ACTL7B-ACTL6A-HSA model in the same position as the INO80 ARP module and added a nucleosome with an extended DNA strand (PDB: 1ZBB) as to exactly emulate the evidence-based structural conformation of the INO80 complex. This diagram assumes that given that SRCAP and INO80 belong to the same protein family then they would form relatively similar structural complexes.

### Loss of ACTL7A and ACTL7B differentially alter lysine acetylation, histone H3, and HDAC localization patterns in murine spermatids

Examining the present data, we have identified the intranuclear presence of ACTL7B (Fig. 1), the activity of a putative NLS (Fig. 2-4), transcriptional changes in the absence of *Actl7b* (Fig. 5-6), and *in silico* models predicting the binding capabilities of both ACTL7A and ACTL7B to four members of prominent nucleosome remodeling families (Fig. 7-9). Altogether, these results indicate involvement of ACTL7A and ACTL7B in chromatin regulation. Given the potential impact of testis specific ARPs contributing to modified chromatin regulatory complex functions, we hypothesized that recruitment of chromatin modifying factors may be impacted. Therefore, we next evaluated evidence for broad changes in nucleosome remodeling activity over the course of spermiogenesis as consequence of ACTL7A or ACTL7B ablation. Given the known importance of histone acetylation as one of the primary epigenetic changes that regulate gene expression, and our previous observation that acetylation in the 7B KO testis was impacted (Ferrer et al, submitted), we focused on evaluating the localization and expression patterns of lysine acetylation, histone H3, and histone deacetylases HDAC1 and HDAC3 across spermatid development among WT, *Actl7a -/-* (7A KO)*, and Actl7b -/-* (7B KO) mice (Fig. 10-11).

**Figure 10:**
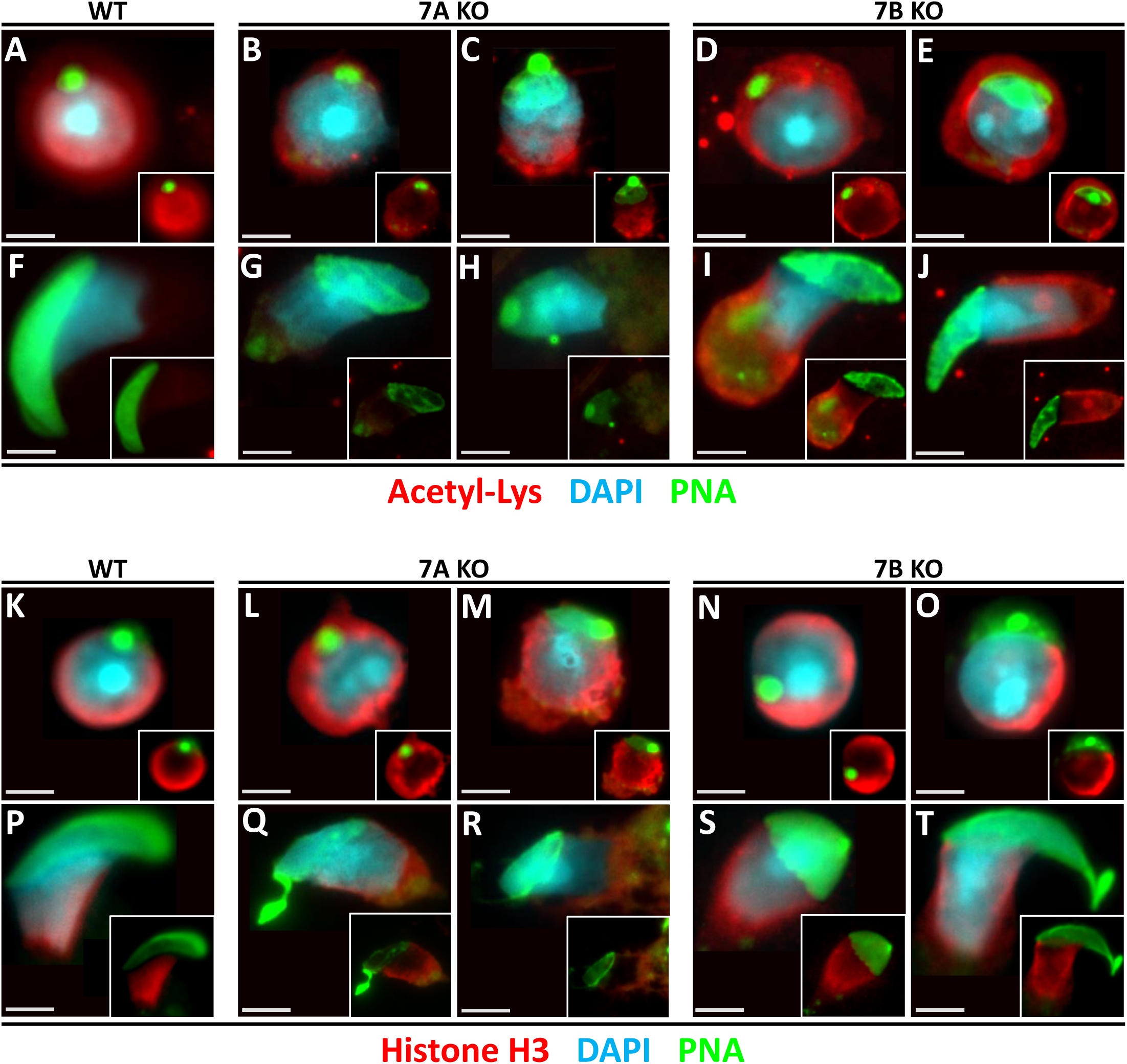
Loss of ACTL7A and ACTL7B Affect Intra-cellular Patterning Changes in Acetylated-Lysine and Histone H3 Across Spermiogenesis. (A-E) Widefield immunofluorescent images of round spermatids showing differential expression patterns of acetylated lysine in their intranuclear spaces with strong nuclear associated expression in WT (A), which is diminished in both *Actl7a -/-* (7A KO) (B-C), and *Actl7b - /-* (7B KO) (D-E) mice. (F-J) Accompanying images of elongating spermatids for each of respective genotype highlighting the reduction of acetylated lysine during these steps of development in both WT and 7AKO, but persistent signal in 7BKO. (K-O) Immunofluorescent images of round spermatids from across the three depicted genotypes illustrating a somewhat conserved expression patterns of histone H3 in their intranuclear spaces. (P-T) Accompanying images of elongating spermatids for each of respective genotypes showing varying degrees of H3 incorporation onto the forming postacrosomal sheath with reduced signal apparent in the nuclear associated postacrosomal sheath of 7AKO. All white scalebars per image represent 2.5 um.

**Figure 11:**
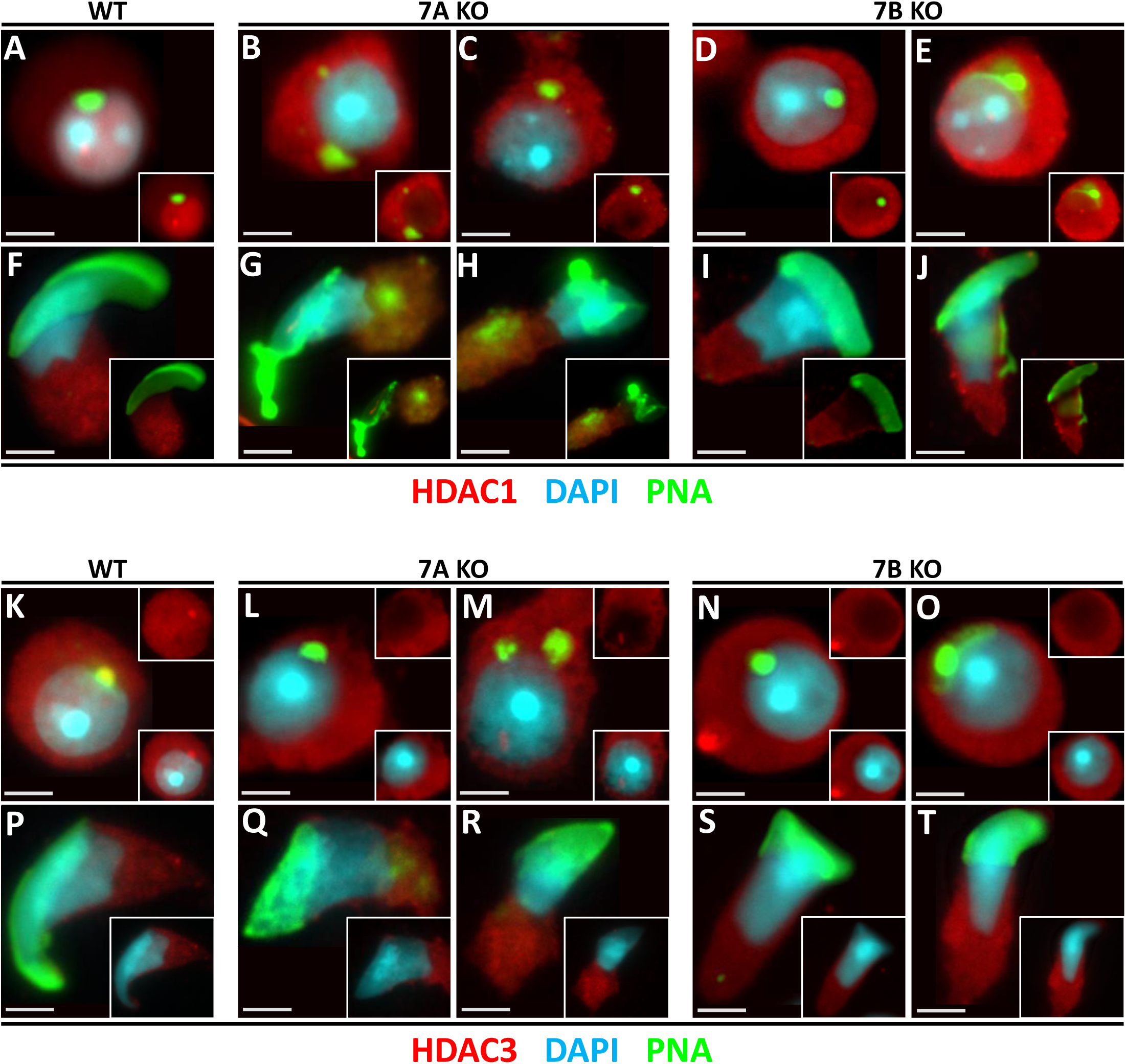
HDAC1 and HDAC3 Localization is Altered within ACTL7A and ACTL7B Null Spermatids. (A-E) Representative widefield immunofluorescent images of round spermatids showing contrasting patterns of HDAC1 expression with WT (A) exhibiting nuclear localization and enhanced expression in a nucleolar associated puncta. *Actl7a -/-* spermatids (7A KO) (B-C) lose these expression patterns and instead exhibit cytoplasm enriched localization, and *Actl7b -/-* (7B KO) (D-E) mice exhibit variable intermediate HDAC1 localization to both cytoplasm and nuclear compartments with reduced observation of nucleolar associated HDAC1 puncta. (F-J) Accompanying images of elongating spermatids for each of respective genotype showing no clear HDAC1 expression differences between them. (K-O) Immunofluorescent images of round spermatids from each genotype illustrating altered intranuclear HDAC3 incorporation. Nuclear localization is diminished in both KOs (L-O) compared to the WT control (K), while cytoplasmic localization is comparable between genotypes. Round spermatids in both KOs also lose acrosomal granule associated HDAC3 localization seen in WT (K). (P-T) Accompanying images of elongating spermatids exhibiting similar cytoplasmic HDAC3 expression patterns between genotypes. All white scalebars per image represent 2.5 um.

Global lysine acetylation in WT spermatids displayed a strong intranuclear signal in round spermatids (Fig. 10A) which was no longer detected in late elongating WT spermatids (Fig. 10F). Importantly, both 7A KO (Fig. 10B-C), and 7B KO (Fig. 10D-E) round spermatids exhibited a loss of intranuclear enrichment of global lysine acetylation detected with this antibody, with the cytoplasmic signal instead exhibiting increased intensity over the intranuclear signal. In elongating spermatids, the 7A KO (Fig. 10G-H) had a lack of acetylated lysine signal akin to the WT control (Fig. 10F) while 7B KO elongating spermatids exhibited a surprisingly strong acetylated lysine signal in their cytoplasmic lobes and around the manchette area indicating aberrant retention (Fig. 10I-J).

Histone H3 localization was comparable between WT, 7A KO, and 7B KO round spermatids (Fig. 10K-O) adopting a strong symmetrical signal at the nuclear periphery with the signal weakening on the nuclear side where the acrosome resides. Previous literature has identified histone H3 to be a component of the postacrosomal sheath beginning during elongation (Hamilton et al., 2021), an observation which we have confirmed in our WT control (Fig. 10P). While the morphology in elongating spermatids in both 7A and 7B makes it difficult to assess the H3 patterns in the caudal aspects of the nucleus, we did note that 7A KO elongating spermatids (Fig. 10Q-R) had difficulty incorporating histone H3 at the caudal aspect of their nuclei where the pattern was concentrated more peripherally. This was evident by the H3 signal-less void delineating the nucleus from the cytoplasm of 7A KO cells (Fig. 10Q-R subpanels) and suggests the residual staining may be cytoplasmic rather than postacrosomal sheath associated in the 7A KO elongating spermatids. In 7B KO elongating spermatids (Fig. 10 S-T) the signals appeared patchy consistent with abnormal elongating spermatid morphology, but these cells still exhibited histone H3 signal across the caudal nuclear surface suggesting a capacity for the expected localization pattern observed in the WT control (Fig. 10P). These postacrosomal observations may not have relevance to nucleosome remodeling activity, however these results may have additional implications regarding the functions of these proteins in the formation and regulation of the postacrosomal sheath.

Focusing on the observed changes in intranuclear lysine acetylation of both 7A KO and 7B KO round spermatids, we sought to elucidate if conventional regulators of acetylated lysine had changed in the KOs. The importance of this undertaking arises from the fact that acetylation of the numerous lysine residues found in the tails of histone octamers is one of the primary ways nucleosome remodelers concomitantly relate to changes in chromatin regulation (Musselman et al., 2012). For this approach, we opted to look at HDAC1 and HDAC3 in our KO models for two reasons. 1) Both HDAC1 and HDAC3 belong to the class I HDAC subfamily which is primarily, if not exclusively, known for robust intranuclear activity (RUIJTER et al., 2003). 2) HDAC1 and HDAC3 are readily expressed across the germline with some known functions. Specifically, HDAC1 has been reported to cooperate with BRDT and possibly PRMT5 and TRIM28 to enact transcriptional repression via their interactions with transcriptional promoters in the germline (Wang & Wolgemuth, 2015). More recently, HDAC3 has been implicated as a key genetic regulator for haploid cell progression and gene regulation in the male germline. In association with NCOR and SOX30, HDAC3 has been exhibited to inhibit the expression of thousands of meiotic/spermatogonial genes while promoting the transcription of post-meiotic haploid genes in equal numbers (Yin et al., 2021). Contextualizing the aforementioned roles of HDAC1 and HDAC3 in spermatogenic chromatin regulation with the changes in intracellular lysine acetylation (Fig. 10A-J), and the altered transcript levels of various transcription factors in 7B KO testis (Fig. 6B-C) led us to hypothesize that alterations to these HDACs might contribute to the detected abnormalities found in the 7A KO and 7B KO germline. Examination of HDAC1 expression in WT round spermatids revealed a strong intranuclear presence with a consistently enriched puncta associated with the periphery of the nucleolus (Fig. 11A). Fascinatingly, all round spermatids observed in 7A and 7B KOs lacked the observed WT pattern of intranuclear HDAC1, instead exhibiting strong cytoplasmic signal (Fig. 11B-E). Additionally, 7A KO round spermatids had a complete loss of the HDAC1 positive peri-nucleolar puncta found in the WT controls (Fig. 11A-C) while 7B KO spermatids retained such HDAC1 peri-nucleolar expression pattern (Fig. 11D-E) in approximately 43% of round spermatids. We observed that localization of the HDAC1 signal shifted in normal WT developmental progression between round and elongating spermatids, with WT late elongating spermatids presenting only a weak cytoplasmic HDAC1 signal (Fig. 11F). Both 7A and 7B KO elongating spermatids also exhibited this pattern suggesting little to no change in elongating spermatid cytoplasmic retention after the aberrant absence of nuclear association in round spermatids (Fig. 11G-J).

HDAC3 localization in WT round spermatids was routinely observed to have an enriched puncta associated with the acrosomal granule, in addition to a ubiquitously homogenous expression pattern between the cytoplasmic and intranuclear space (Fig. 11K). Interestingly, 7A and 7B KO round spermatids both exhibited a loss of HDAC3 signal in both the intranuclear space and the acrosome granule (Fig. 11L-O). Elongating spermatids across all three genotypes exhibited the same cytoplasmic expression pattern of HDAC3 with no nuclear presence nor associations with the now flattened acrosomal granule (Fig. 11P-T). Altogether, the observed changes in lysine acetylation and HDAC localization reflect a pronounced shift in nuclear dynamics of these chromatin regulatory associated signals in round spermatids.

## Discussion

We here proposed a novel mechanistic role for testis-specific actin related proteins ACTL7A and ACTL7B in chromatin regulation during spermiogenesis and male fertility through contribution to chromatin regulatory complexes known to coordinate epigenetic nuclear dynamics. The proposed nuclear roles are supported by combined *in vivo*, *in vitro*, and *in silico* evidence. Our proposed mechanistic model builds from known nuclear roles for somatic ARPs that have established functional and structural models.

Utilizing these models, we derived an AI facilitated *in silico* approach to model ARP subunit swapping with testis-specific ACTL7A and ACTL7B utilizing AlphaFold2. We have shown *in silico* evidence for the reliability of AlphaFold2 to predict the interaction between ARPs and HSA domains and revealed the potential for ACTL7A and ACTL7B to directly participate in nucleosome remodeling activities through the HSA domain of prominent chromatin regulators such as the INO80 and SWI/SNF complexes. We further show that the predicted associations with various combinations of ARPs have differential predicted biological relevance scores and can adopt different conformations. These interaction models thus suggest the potential for spaciotemporal co-localizations and stoichiometric ratios of testis expressed actins and ARPs to impact the specific composition and function of nucleosome remodeling complexes impacting germline chromatin regulation throughout spermiogenesis.

Potential regulatory implications for predicted testis specific chromatin remodelers may be derived from examples of such roles supported by evidence in somatic INO80 and SWI/SNF complexes. For example, the INO80 family complexes execute nucleosome sliding and aid in DNA replication through the canonical INO80 complex in yeast and hINO80 complex in humans (Knoll et al., 2018; Zhang et al., 2023). Additionally, CREB and NOTCH-mediated transcriptional coactivation as well as DNA repair through the exchange of histone variant H2A.Z for canonical H2A in nucleosomes is mediated by INO80 family member SWR1 in yeast and SRCAP in humans (Eissenberg et al., 2005; Colino-Sanguino et al., 2021). INO80 family complexes are known to include ARP4, ARP6, ARP8, and actin in yeast and ACTL6A/BAF53B (ARP4 ortholog), ACTL6B/BAF53B (ARP4 ortholog), ACTR6 (ARP6 ortholog), ACTR8 (ARP8 ortholog) and β-actin in mammals (Knoll et al., 2018; Klages-Mundt et al., 2018). Additionally, both INO80 and SWR1/SRCAP complexes use a RuvB1/2 (RuvBL1/2 in mammals) hexamer as a scaffolding cofactor for their nucleosome regulating activity, a dependency which is highly conserved in this family of nucleosome remodelers (Willhoft & Wigley, 2020). In contrast, SWI/SNF complexes do not require an association with a RuvB hexamer and are implicated in transcriptional repression and upregulation through ZEB1 and CREST-dependent pathways (Sánchez-Tilló et al., 2010; Qiu & Ghosh, 2008) as well as have been characterized to be crucial components for the differentiation and development of cellular lineages (Paula Coutinho Toto et al., 2016; Alver et al., 2017; Smith et al., 2022). SWI/SNF complexes are known to contain Arp4, Arp6, Arp7, Arp8, Arp9, and actin in yeast while the mammalian equivalent BAF complexes contain ACTL6A, ACTL6B, and β-actin (Bieluszewski et al., 2023). Thus, we speculate that testis-specific ARP subunit swapping in chromatin remodelers may lead to germline-specific modification of processes such as affinity based associations, recruitment of epigenetic regulators, association with specific histone variants, nucleosome PTM recognition, and/or direct DNA binding. Our proposed mechanism of testis-specific ARP contributions to testis-specific chromatin remodeling by contributing to HSA domain containing chromatin remodelers through ARP subunit swapping forecasts a plethora of future work to validate potential interactions and investigate potential implications of such putative complexes implicated by our predicated modeling.

As a first step in establishing *in situ* evidence for the potential of ACTL7A and ACTL7B to contribute to nuclear regulation, we also provide here evidence of nuclear localization and trafficking regulation, transcriptional consequences of targeted ablation in mouse models, and ACTL7A and ACTL7B dependent nuclear association of regulatory HDACs. This builds on our previous work demonstrating that these two ARPs are both separately causative of male infertility in KO mouse models exhibiting ultrastructural defects to the actin-associated acrosomal cytoskeleton (Ferrer et al., 2023) and the flagellum (Clement et al., 2023). Given the diverse and at times unexpected roles that many ARPs have been observed to have, we hypothesized that ACTL7A and ACTL7B may functionally play roles beyond observed structural and cytoplasmic roles in spermiogenesis. Specifically, we proposed these ARPs as key molecular components of spermiogenesis through additional nuclear regulatory contributions. In this study we began by revisiting nuclear localization of testis specific ARPs. Our previous characterization of ACTL7B localization in the germline utilized commercial antibodies derived from immunogen sequences containing large portions of the actin body of ACTL7B, a protein domain in which ACTL7B largely shares homology with other actins (Chadwick et al., 1999; Clement et al., 2023). In contrast, the immunogen sequence that derived the antibody used in this study was from the unique N-terminal sequence of ACTL7B. With this N-terminal specific antibody we reaffirmed our previous finding of ACTL7B being associated with the acrosomal region, and discerned its presence in the nucleoplasm of round spermatids. The previous antibody used detected ACTL7B to be present in nuclei only in round spermatids. However, with the N-terminal specific antibody, we consistently observed ACTL7B specific positive signals emanating from the nuclei of spermatocytes in a pattern consistent with synapsed chromosomes (Qiao et al., 2012). While it seems unlikely that ACTL7B would be a core component of the synaptonemal complex given the obvious lack of meiotic checkpoint failure in the *Actl7b -/-* mice, it is possible that ACTL7B plays a role in the transcriptional priming of haploid germ-cell expressed genes found in the chromatin loops protruding from the synaptonemal complex in pachynema (Prakash et al., 2015). Further investigation is required to determine the significance of this spaciotemporal localization of ACTL7B within spermatocyte nuclei.

In addition to the presence of ACTL7B in the nucleoplasm, we have identified the actin-like domain of this protein to be responsible for its nuclear affinity rather than its specialized N-terminal domain (Fig. 3). More specifically, we have identified a highly conserved tri-lysine repeat in the fourth actin subdomain to be capable of the transposition of ACTL7B from the cytoplasm to the intranuclear space.

We cautiously present this as a potential directly encoded NLS for testis-specific ACTL7B given that most actins and ARPs are conventionally known to be transported to the nucleus by forming a tetrameric complex with cofilin and an importin α/β dimer, where the associated importins recognize an NLS within cofilin rather than directly binding the actin (Kristó et al., 2016). Despite the tripartite interaction between actins, cofilin, and importins, there is an emergent body of literature that has suggestive evidence of some ARPs not relying on cofilin for their nuclear localization (Kristó et al., 2016; Kelpsch & Tootle, 2018). Moreover, the evidence for ACTL7B having a directly encoded NLS is also further supported by the presence of importins and the absence of cofilin in co-purified fractions using ACTL7B as bait and analyzed through mass spectrometry (Ferrer *et al*. Submitted). Nevertheless, despite the somewhat conflicting evidence of ARPs having ulterior methods of nuclear transport, a topic that is still debated and periodically reexamined, we present the possibility of ACTL7B containing a directly encoded NLS which may offer alternative or additional means of spaciotemporal dynamics in spermatogenesis.

After affirming the presence of ACTL7B in the nucleus, we investigated the transcriptomic consequences of ACTL7B ablation as an initial query of potential involvement in chromatin regulation. Initial analysis of the transcriptomic data showed that a vast majority of altered transcripts were upregulated in *Actl7b* -/- testes. Most of the upregulated transcripts belong to known molecular pathways for the regulation of protease inhibition, cell death, the extracellular matrix, and immune/inflammation responses. This initial screening of the transcriptome was quite unexpected, though with further reflection we believe to have an explanatory hypothesis for these observations. We have observed that ACTL7B is present in the subacrosomal region of spermatids (Clement et al., 2023;Ferrer et al. submitted), and that other testis-specific actins like ACTL7A (Ferrer et al., 2023) and ACTRT1 (Zhang et al., 2022) have also been shown to be structural components of the subacrosomal space. Individual ablations of these three testis-specific actins have also shown a conserved phenotype of acrosomal detachment or “peeling acrosomes”. It stands to reason that in these cases where acrosomal integrity is disrupted during spermiogenesis the peeled acrosomes may expose the surrounding seminiferous epithelium to the acidic and enzyme-rich acrosomal components. This increase in enzymatic exposure could explain the transcriptional surge of irreversible protease inhibitors in order to properly neutralize acrosomal contents (Fig. 5). Furthermore, given the caustic nature of the acrosomal composition (Aldana et al., 2021), it is also possible that chronic exposure to peeling acrosomes in *Actl7b* -/- testes could lead to irritation or inflammation of the seminiferous epithelium. Often times, inflammatory responses lead to immune recruitment (Chen et al., 2018) which both in part require dissolution of the extracellular matrix for cellular infiltration (Sutherland et al., 2023), as well as additional signaling cascades that incur cell death like the complement system (Martin & Blom, 2016). Notably, each of these systems had many associated components highly upregulated in the observed *Actl7b* -/- testes transcriptome (Fig. 5). Further evidence is clearly needed to support these implications. However, if the hypothetical courses of events caused by acrosomal instability were found true, it could affect a significant number of men with acrosomal defects and be categorized as a novel contributor to male infertility. Many human polymorphisms have been recently reported of single-point coding mutation in subacrosomal proteins like ACTL7A, ACTL7C, and ACTRT1 (Xin et al., 2020; Wei, Wang, et al., 2023; Zhou et al., 2023; Sha et al., 2021) that lead to both male infertility and peeling acrosomes. If the presented transcription changes in *Actl7b* -/- testes were caused in part by the exposure to acrosomal contents leading to accompanied inflammation, immune, and apoptotic responses, then a similar hypothesized pathology would be expected in men with these polymorphisms. We propose that this could be described as an “Acrid Testis Syndrome” where the prerequisite of testicular acrosomal content exposure could lead to the described pathology.

Looking at additional transcriptional alterations in *Actl7b* -/- testes, we noted a significant change in several transcription factors, some of which are known to be specifically enriched at different developmental stages of the male germline (Green et al., 2018). When looking at these germline specific transcription factors, loss of ACTL7B led to an increase in spermatogonia transcription factors while significantly reducing spermatid specific transcription factors. This curious observation was also previously observed in HDAC3 deficient mice (Yin et al., 2021), which incurs further interest given the lack of intranuclear HDAC3 presence in both ACTL7A and ACTL7B deficient spermatids (Fig. 11) suggesting that altered germ-line transcription factor levels downstream of ACTL7B ablation may be mediated through HDAC3. If HDAC3 fails to be properly incorporated in the nuclear space of the germline given the loss of these ARPs, then in theory it could not enforce its previously described roles as a premeiotic gene repressor and haploid gene enhancer with SOX30 and NCOR (Yin et al., 2021). Nuclear actins have previously been shown to monomerically bind and regulate the activity of class I HDACs, specifically HDAC1 and HDAC2 (Serebryannyy et al., 2016). This, however, would not directly explain how the absence of specific ARPs would affect the lack of intranuclear class I HDAC signals as we have observed (Fig. 11). Three possible hypotheses are: 1) ACTL7A and/or ACTL7B are required to recruit, guide, or incorporate HDACs 1 and 3 into chromatin regulating complexes in a haploid-specific context. In the absence of these ARPs, affected HDACs would thus be left derelict in the nucleoplasm as monomeric subunits and become more susceptible to nuclear export through now exposed molecular motifs like nuclear export sequences. 2) In the absence of ACTL7A or ACTL7B other ARPs bind these HDACs, in the same inhibitory fashion as with nuclear actin and HDAC 1 (Serebryannyy et al., 2016), resulting in the occlusion of the epitope recognized by the antibody used in this study. 3) Through a currently unknown mechanism ACTL7A or ACTL7B co-chaperone HDACs 1 and 3 during their nuclear transport or indirectly affect the phosphorylation state of their NLS, in a similar fashion as Class IIa HDACs are regulated (Mathias et al., 2015), thus modulating their nuclear transport. Relatedly, the similarities observed in this study in the irregular localization of HDACs 1 and 3, as well as the loss of intranuclear lysine acetylation in both ACTL7A and ACTL7B deficient round spermatids may be indicative of direct roles for both ARPs, however given the transcriptomic and protein level evidence presented here showing that loss of ACTL7B leads to a severe downregulation of ACTL7A, we hypothesize that *Actl7b* -/- mice can phenotypically be considered as an ACTL7A knock-down (KD) model. Thus, the more sever phenotype and any individual effects overlapping with *Actl7a* KOs may be interpreted as the result of the deficiency of one ARP or both, while clearly implicating ACTL7B in the regulation of *Actl7a* expression. In either case, we propose that these testis specific ARPs regulate HDAC localization with a paradoxical effect on loss of both nuclear HDACs and intranuclear acetylation in ARP KOs which warrants future mechanistic investigation.

In summary, this study has investigated the intranuclear functionality of both ACTL7A and ACTL7B. We have reported affirmation of the intranuclear presence of ACTL7B, discovered a strong candidate for its nuclear localization sequence, identified the gross transcriptomic consequences of ACTL7B ablation, shown evidence of the transcriptional dependence of ACTL7A to the presence of ACTL7B, and identified changes in HDAC expression patterns and lysine acetylation in both *Actl7a* -/- and *Actl7b* -/- spermatids. Furthermore, we have used an AI facilitated *in silico* modeling approach to provide evidence for the potential association of testis-specific ARPs with canonical somatic chromatin regulatory complexes containing HSA domains and propose a mechanistic role for testis-specific ARPs in subunit swapping within these complexes for unique contributions to spermiogenic chromatin dynamics. Together with our previous observations of cytoplasmic and cytoskeletal structural roles, we propose that ACTL7A and ACTL7B serve as spaciotemporal coordinators of spermiogenesis through multiple intracellular mechanisms related the known functions of somatic ARPs. This is an exciting prospect as somatic ARPs tend to specialize, suggesting that these testis ARPs may have evolved to facilitate coordination of the characteristically unique temporally associated restructuring of diverse cellular components during spermiogenesis including structural cytoskeletal and nuclear regulators. Ultimately, these findings further our understanding of the indispensable roles of ACTL7A and ACTL7B in male fertility as well as our understanding of the broader roles for the ARP family of proteins.

## Materials and Methods

### Ethics statement

All experiments involving animals were approved by the Institutional Animal Care and Use Committees of Texas A&M University (IACUC #2018-0104) and were conducted in compliance with the guidelines and regulations for animal experimentation of this institution.

### Generation of Actl7a and Actl7b KO mice

*Actl7a* -/- (7A KO) mice were produced through the application of CRISPR/Cas9 technology at Osaka University as previously reported (Abbasi et al., 2018, Ferrer et al. 2023). The *Actl7b* -/- (7B KO) mouse line was produced by injecting modified ESCs into blastocysts to generate chimeric founders and selecting for germ-line transmission to establish the line as previously reported (Clement et at 2023). Mouse colonies were established at Texas A&M University, were housed in a 12-hour light and 12-hour dark diurnal cycle and were provided unrestricted access to food and water.

### Plasmid expression and HEK239F transfection

The YFP-tagged full-length human ACTL7B vector and various truncated ACTL7B vectors, were constructed using the Gateway cloning system (Invitrogen, Waltham, MA, USA). The process entailed cloning the various human ACTL7B gene variants into the pENTR TOPO vectors using the pENTR d-Topo cloning kit (Invitrogen), following the manufacturer’s guidelines. The full-length ACTL7B expression vector as well as its truncated variants were generated through LR clonase reaction (Gateway® LR clonase™ II Enzyme Mix, Invitrogen, Waltham, MA, USA) using the destination vector pcDNA6.2/N-YFP-DEST (Supplementary Fig. S1A-F). To serve as experimental controls, complementary vectors were constructed lacking the ACTL7B coding sequence or including the SV40 NLS [PKKKRKV] (Kalderon et al., 1984). Sequence integrity of vectors were validated through DNA sequencing. The Top10 chemically competent Escherichia coli cells were used for cloning and plasmid amplification. The Endo-free plasmid maxi kit (#12162, Qiagen, Valencia, CA, USA) was used for plasmid purification.

HEK293F cells (#R790-07, ThermoFisher Scientific, Waltham, MA, USA) were cultured and transfected with the pcDNA6.2N-YFP-ACTL7B plasmids using the Freestyle 293 Expression System. In brief, HEK293F cells were maintained in suspension within Freestyle™ 293 Expression Medium (#12338026, ThermoFisher Scientific, Waltham, MA, USA) at a temperature of 37°C and 5% CO2. Passage of the cells occurred when their population density ranged between 1 × 10E6 and 3 × 10E6 cells per ml of media proven that their cell viability exceeded 94%. Following approximately three passages, the HEK293F cell culture underwent transfection with a ratio of 1 part pcDNA6.2N plasmid to three parts 293Fectin (#12347019, ThermoFisher Scientific, Waltham, MA, USA). After transfection, the cells were cultivated for a duration of 17-24 h, then fixed using 4% PFA for a period of 1 h, after which they were stored at 4°C for subsequent immunofluorescence evaluations.

### Germ Cell dissociation and Immunofluorescent microscopy

Single cell suspensions were prepared as previously described (Ferrer et al. 2023). Briefly, adult murine testes with their tunica albuginea removed were placed in 10 ml of Krebs-Ringer bicarbonate buffer (EKRB) (Bellve et al. in 1977) and subjected to enzymatic digestion with 2.5 mg of collagenase IV (#SCR103, Sigma-Aldrich, Saint Louis, MO, USA) for 15 min within a shaking incubator at 37°C, 5% CO2, and 150 rpm. This was followed by a treatment involving 5 mg of trypsin III from bovine pancreas (#T9201, Sigma, Saint Louis, MO, USA) along with 1 mg of DNase I (#D4527, Sigma, Saint Louis, MO, USA) in 10ml EKRB for a duration of 30 min. The digested tissue was then homogenized through gentle pipetting for 4 min, after which 5 mg of trypsin inhibitor (#T9003, Sigma-Aldrich, Saint Louis, MO, USA) was introduced upon completion of this procedure. Lastly, the resulting singularized cell mixture was subjected to gravity filtration through a 70 µm nylon mesh (#352350, Falcon, Cary, NC, USA), followed by centrifugation at 250×g for 10 min, and fixation with 4% PFA for a duration of 30–60 min.

For immunofluorescent microscopy, isolated germ cells or transfected HEK293F cells were settled overnight onto slides coated with poly-L-lysine. The settled slides were then subjected to permeabilization using PBS containing 1% Triton X-100 (PBST) for a duration of 15 min. Slides were thoroughly washed with PBS and blocked with 10% horse serum in PBS for 30 min. The staining process involved incubating the slides with rabbit and/or mouse primary antibodies, at their respective dilutions (Supplementary Table S1), in PBS for a duration of 2 h. Following primary antibody incubation, slides were washed thrice with PBS for 5 min per wash, then incubated with Alexa-conjugated secondary antibodies for a duration of 1 h. When performing co-labeling with fluorescent conjugated peanut agglutinin (PNA), a dilution of 1:400 was applied for a duration of 1 h concurrent with the secondary antibody incubation. For nuclear counter-staining, DAPI was used at a dilution of 1:500 for a duration of 3 min. Slides were mounted using Aqua-Polymount (#18606, Polysciences, Warrington, PA, USA) and imaged using either a Leica Dmi8 S Platform (LAS X software V5.0.2) inverted microscope system for widefield microscopy, or a Zeiss LSM 780 multiphoton microscope (ZEN Blue software V3.4) for confocal microscopy. The resulting images of germ cells and HEK293F cells were analyzed to elucidate patterns of localization relative to the specific cell type and, in the case of germ cells, their developmental stage. A comprehensive listing of all antibodies employed can be found in (Supplementary Table S1). All intracellular localization assessments of protein targets via immunofluorescence were performed in triplicates with cells extracted from 3 separate mice per genotype in question (WT, ACTL7A KO, and/or ACTL7B KO). For specific spermiogenic developmental assessments of protein targets (Histone H3, Acetylated Lysine, HDAC1, and HDAC3), 30-50 cells per mouse per genotype were assessed separately for both round and elongating spermatids.

### Transcriptomic processing and analysis

Testes from three *Actl7b* -/- and three littermate WT mice were collected and snap frozen in liquid nitrogen before RNA isolation using TRIzol reagent according to the manufacturers protocol (Invitrogen). RNA quality was assessed by Agilent TapeStation 4200 and analysis software version 3.2. RNA samples were diluted to 75ng/ul for a total of 750ng RNA total input for library generation. RNA libraries were prepared using the TruSeq Stranded Total RNA method according to the manufacturers instructions (Illumina). Qubit values were taken of the final library and the nM concentration was calculated based on the concentration and base pair size with initial library concentrations ranging from 33.7-40.5 ng/ul.

Libraries were then diluted to 4nM concentration for sequencing on an Illumina NovaSeq 6000 platform. The sequencing data was processed and analyzed by using the public Galaxy server environment (Afgan et al., 2022). Adapter sequences, low quality, and short reads were removed using Trimmomatic (Bolger et al., 2014). Trimmed reads were then mapped using HISAT2 [PMID: 31375807] with the mm10 (Mus musculus) reference genome. The number of reads per annotated gene was counted using featureCounts (Liao et al., 2013). Differential expression testing was conducted using DESeq2 (Love et al., 2014). For further analysis and data visualization we used SRPlots to generate the present volcano plot and GO pathway analysis charts (http://www.bioinformatics.com.cn/), TFCheckpoint2 for detecting transcription factors in the population of altered transcripts (Chawla et al., 2013), GraphPad Prism version 9.0 and Single-Cell RNA-Seq data from (Green et al., 2018) to detect and visualize changes in germ cell specific transcription factors, and MORPHEUS (https://software.broadinstitute.org/morpheus) for all generated heatmaps and accompanied clustering analysis.

### *In silico* AlphaFold 2 multimer interaction modeling

All AlphaFold 2 predictions exclusively used the human orthologues of each protein mentioned. Protein interaction models between all components were performed in accordance with the previously described methodologies for elucidating multimeric protein complexes (Jumper et al., 2021; Evans et al., 2022). In short, all multimeric protein models were performed through AlphaFold 2 inside a Linux virtual machine accessed through the Windows Subsystem for Linux (WSL) application. The windows workstation used was equipped with a Ryzen 5800X CPU, RTX 3090 24 gigabytes vRAM GPU, and 64 gigabytes of DDR4 RAM to satisfy the high hardware demands of the application. With the 24 gigabytes of vRAM available we were able to comfortably run protein complex prediction up to 1750 amino acid residues in size. All protein models were also performed with GPU relaxation enabled, three recycles per neural network, and with templates and amber disabled to expedite runtime while maximizing model fidelity (Evans et al., 2021). For structural assessments, all multimeric protein models were visualized and analyzed using ChimeraX V1.4 (Goddard et al., 2017; Pettersen et al., 2020). Predicted alignment error heatmaps and interface predicted template modeling (ipTM) scores were extracted from the AlphaFold predictions and weighed in context with residue interaction, electrostatic potential/hydrophobicity surface, and interface analysis tools in ChimeraX to validate the biochemical feasibility of the predicted protein interactions. Once structural models were validated, the ipTM predicted endpoint was used to numerically represent the stability (i.e., biochemical feasibility) of the predicted interactions. An ipTM score of 0.7 out of 1.0 or higher for dimeric complexes, and an ipTM score 0.6 out of 1.0 or higher for tetrameric complexes, given the lower stability of larger interactions, were considered to be biologically relevant as contextualized by previous studies (Yin et al., 2022; Evans et al., 2022).

### Western blotting

Testicular and sperm samples were suspended in a low salt buffer solution composed of 10 mM Tris-HCl, 205 µM CaCl2, and 100 mM KCl, with a pH of 7.8 before further processing with a Dounce homogenizer. The resulting homogenized lysate mixtures were centrifuged at 10,000×g for 15 min at 4°C. This step aimed to facilitate the collection of the supernatant, constituting the soluble fraction, which was intended for western blot analysis. Following the centrifugation, each lysate was diluted at a 1:1 ratio with a solution of 2× Laemmli sample buffer (#1610737EDU, Bio-Rad, Hercules, CA, USA), containing 5% (v/v) anhydrous 2-mercaptoethanol (#1610710XTU, Bio-Rad, Hercules, CA, USA). The resulting mixture was then boiled for 15 min and then 35 µl of each sample was loaded into individual wells of 10–20%, Tris-Glycine, 1.0 mm denaturing gels (#XP10200BOX, Invitrogen, Waltham, MA, USA). After all protein constituents were separated by gel electrophoresis, they were transferred onto a polyvinylidene difluoride membrane (#1620174, Bio-Rad, Hercules, CA, USA). Following the transfer, the membrane underwent washing with TBST (0.1% Tween-20) and was blocked overnight at 4°C, with a 5% dry nonfat milk solution. Following the overnight block, membranes were incubated with a rabbit anti-ACTL7A primary antibody (Supplementary Table S1) at a dilution factor of 1:5000 for of 2 hours with a 5% milk content in TBST. Corresponding secondary antibodies, specifically anti-rabbit conjugated with horseradish peroxidase, were introduced at a dilution factor of 1:10 000 in TBST. Lastly, the resulting western blot was then imaged using the Bio-Rad ChemiDoc XRS Imaging System and analyzed via the ImageLabs software for band intensity and specificity.

## Acknowledgments

We would like to thank Dr. Mike Godling (Department of Veterinary Pharmacology and Physiology, School of Veterinary Medicine Texas A&M University) for providing aliquots of the histone H3, Acetylated lysine, HDAC1, and HDAC3 antibodies used in this study.

## Authors roles

Conception and design of the study: PF and TMC, Vector production and cell transfection: SU, TMC, and PF, Experimental design, and data acquisition: PF and TMC, analysis and interpretation of data: PF and TMC, Transcriptomic data processing: JJC, PF. Drafting of the article: PF. Critical revision and approval of the final draft and agreement for accountability and integrity of the work: PF, SU, JJC, TMC.

## Funding

This research was supported by the Eunice Kennedy Shriver National Institute of Child Health and Human Development of the National Institutes of Health (NIH R00 HD081204 to Dr. Clement), and by start-up funds to T.M. Clement from the Department of Veterinary Physiology and Pharmacology at Texas A&M University. Pierre Ferrer was supported in part through the NIH training grant T32 ES026568.

The content is solely the responsibility of the authors and does not necessarily represent the official views of the National Institutes of Health.

## Conflict of interest

The authors have no conflicts to disclose.

## Supplemental Figures

**Supplemental Fig 1:**
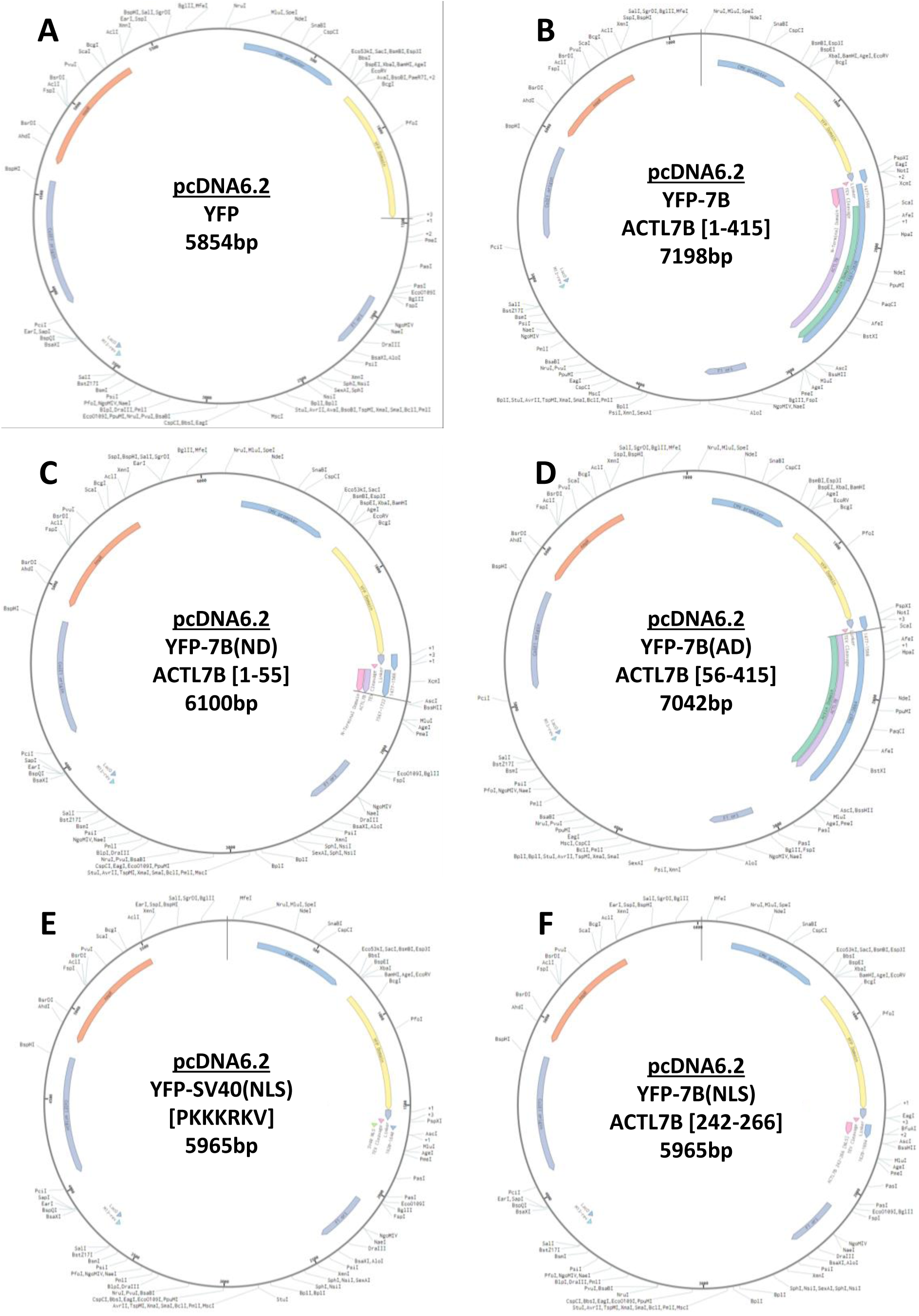
Plasmids. A) YFP only negative control. B) YFP conjugated full length human ACTL7B. C) YFP conjugated N-terminus of human ACTL7B. D) YFP conjugated actin body of human ACTL7B. E) YFP conjugated SV40 NLS positive control for nuclear localization. F) YFP conjugated NLS candidate 2 of human ACTL7B [242-266].

**Supplemental Fig 2:**
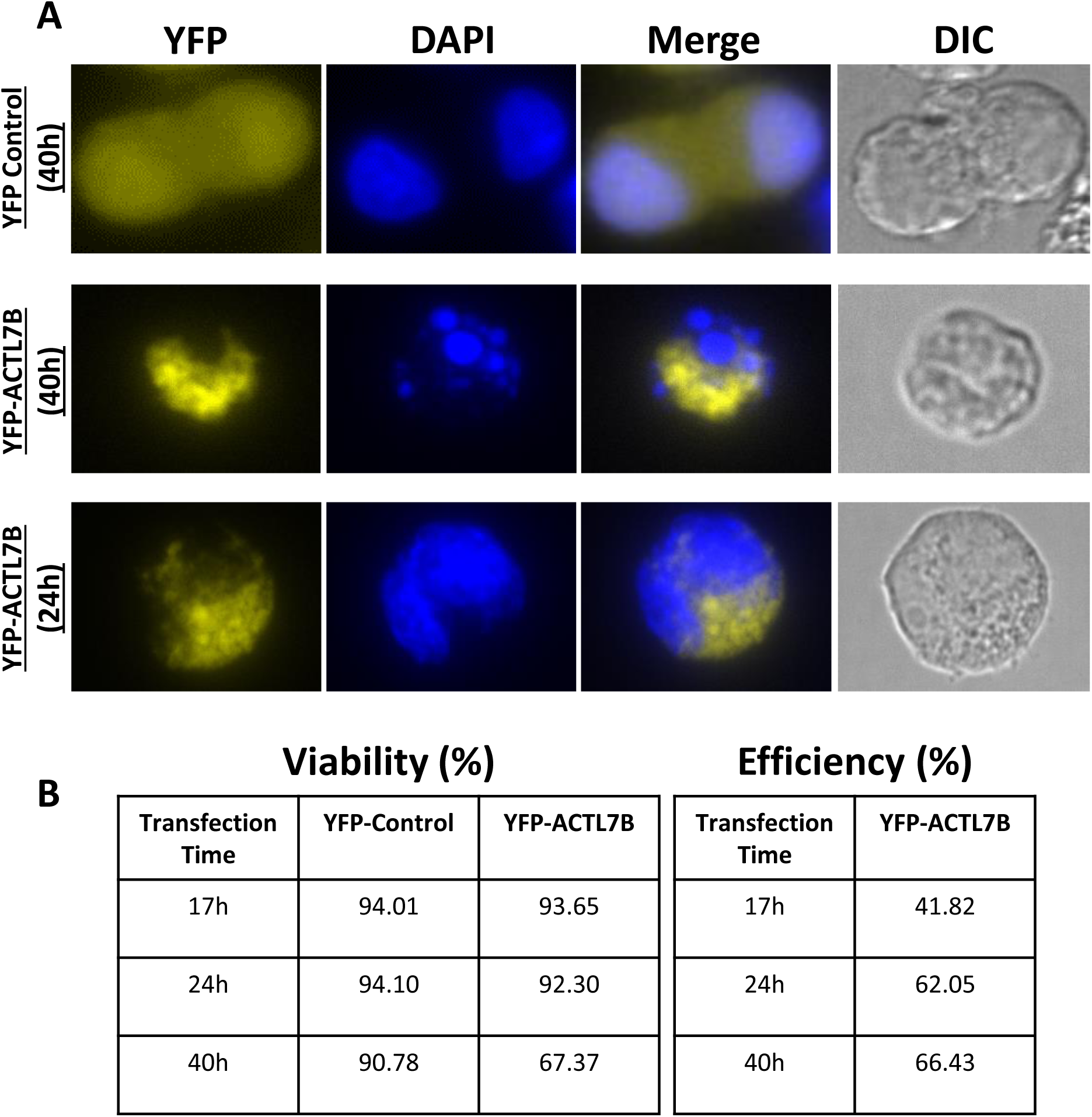
Transfection optimization. A) fluorescent microscopy showing YFP localization patters when conjugated to ACTL7B 24 to 40 hours post transfection. The 40 hours incubation time after YFP-ACTL7B transfection readily led to cell death. B) Tabulated results of cell viability and transfection efficiency indicating a clear decrease in cell viability in YFP-ACTL7B transfected cells compared to the YFP control after 40 hours, but not 17 or 24 hours.

**Supplemental Fig 3:**
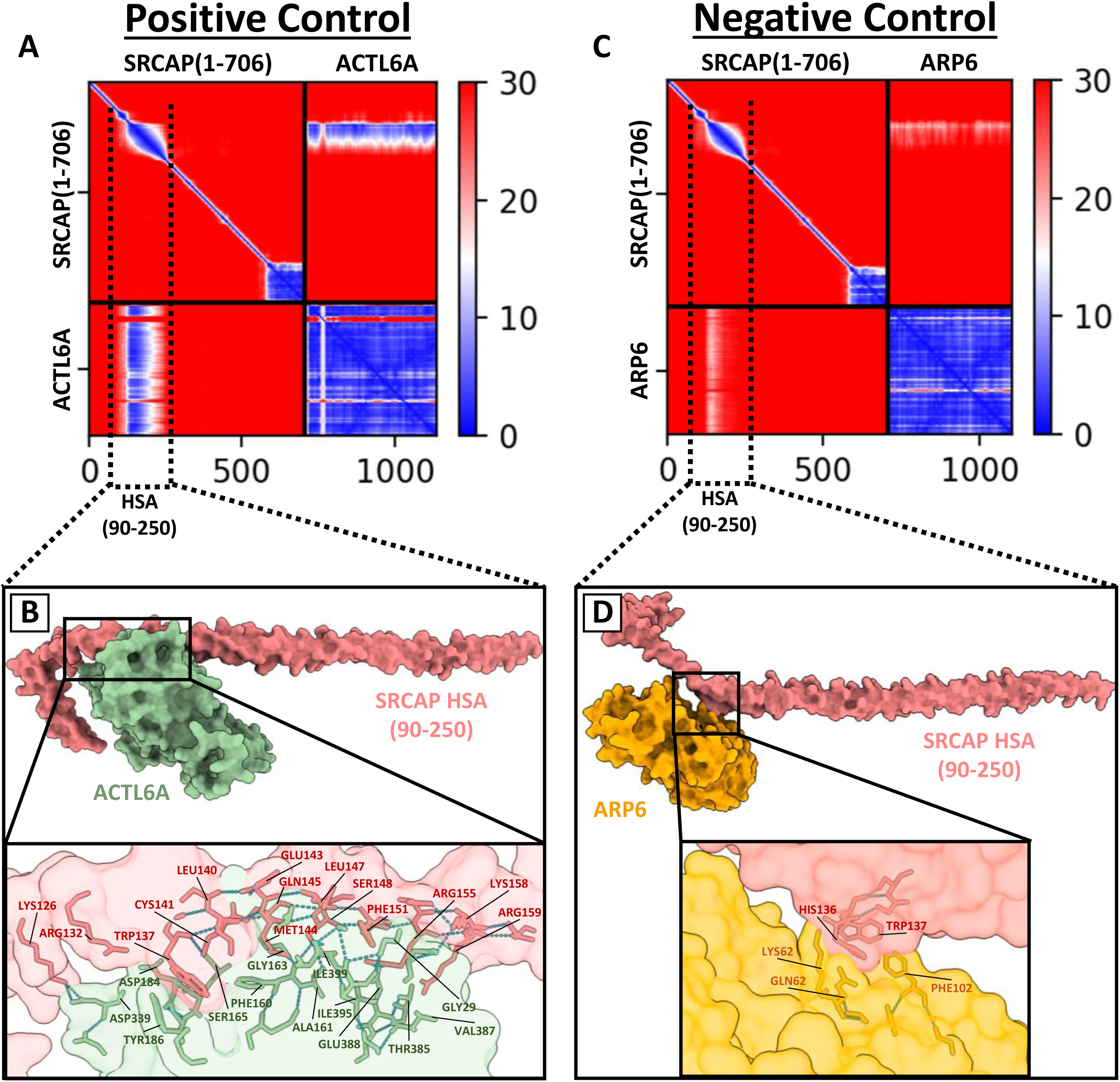
SRCAP HSA-ARP modeling controls. A) Predicted alignment error heatmap of interactions between SRCAP(1-706) and ACTL6A serving as a positive control for a known interaction. Red indicates poor to no binding, while blue on the heatmap indicates excellent binding between the two proteins. The presence of a clear blue vertical strip between the interaction quadrant of ACTL6A with SRCAP indicates the placement of their interaction allowing identification of the HSA domain. B) Generated PDB file of the predicted interaction between ACTL6A and the identified HSA domain of SRCAP. Subpanel illustrates a transparent cross section of the protein-protein interaction showing a vast number of buried (interacting) amino acid residues that stabilize the binding of ACTL6A with SRCAP and associated H-bonds shown as blue dash lines (buried/interacting residues were analytically identified by the interface mapping function in ChimeraX). C-D) Predictive interaction between SRCAP(1-706) and ARP6 serving as a negative control given that ARP6 is known to not interact with SRCAP in this region of the protein. Cross sectional protein mapping of the amino acid residues that interact between the HSA domain of SRCAP and ARP6 show minimal cooperative binding.

**Supplemental Table 1:** Antibodies. Comprehensive list of all used antibodies and other labels; key experimental parameters are indicated.

| <u>Antibody</u> | <u>Manufacturer</u> | <u>Use & Working Dilution</u> |
| --- | --- | --- |
| Alexa 568 Lectin-PNA | Invitrogen, # L32458 | Fluorescent Microscopy (1:400) |
| Alexa 488 Donkey anti-Rabbit | Invitrogen, # A21206 | Fluorescent Microscopy (1:500) |
| Alexa 488 Goat anti-Mouse | Invitrogen, # | Fluorescent Microscopy (1:500) |
| HRP Goat anti-Rabbit | PerkinElmer, # NEF812001EA | Western Blot (1:5000) |
| Rabbit anti-ACTL7A | ProteinTech, # 17355 | Western Blot (1:5000) |
| Rabbit anti-ACTL7B | In house (Clement et al., 2023) | Fluorescent Microscopy (1:500) |
| RM anti-Acetylated Lysine | ProteinTech, # 66289 | Fluorescent Microscopy (1:200)<br>*Citrate Antigen Retrieval 10 min |
| RM anti-Histone H3 | Abcam, # ab1791 | Fluorescent Microscopy (1:150)<br>*Citrate Antigen Retrieval 10 min |
| RM anti-HDAC1 | Abcam, # ab7028 | Fluorescent Microscopy (1:100)<br>*Citrate Antigen Retrieval 10 min |
| RM anti-HDAC3 | Abcam, # ab7030 | Fluorescent Microscopy (1:100)<br>*Citrate Antigen Retrieval 10 min |

## Notes

### Competing Interest Statement

The authors have declared no competing interest.

